# *BirthSeq*, a new method to isolate and analyze dated cells from any tissue in vertebrates

**DOI:** 10.1101/2023.10.10.559090

**Authors:** Eneritz Rueda-Alaña, Marco Grillo, Enrique Vazquez, Sergio Marco Salas, Rodrigo Senovilla-Ganzo, Laura Escobar, Ana Quintas, Alberto Benguría, Ana María Aransay, Ana Dopazo, Juan Manuel Encinas, Mats Nilsson, Fernando García-Moreno

## Abstract

Embryonic development is a complex and dynamic process that unfolds over time and involves the production of increasing numbers of cells, as well as the diversification of different cell types. The impact of developmental time on the formation of the central nervous system is well-documented, with evidence showing that time plays a critical role in establishing the identity of neuronal subtypes. However, the study of how time translates into genetic instructions driving cell fate is limited by the scarcity of suitable experimental tools. We introduce *BirthSeq*, a new method for isolating and analyzing cells based on their birth date. This innovative technique allows for *in vivo* labeling of cells, isolation via FACS, and analysis using high-throughput techniques. We demonstrate the effectiveness of BirthSeq for single-cell RNA sequencing and novel spatially resolved transcriptomic approaches in brain development across three vertebrate species (mouse, chick, and gecko). Overall, BirthSeq provides a versatile tool for studying any tissue in any vertebrate organism, helping to fill the necessity in developmental biology research by targeting cells and their temporal cues.

**SUMMARY STATEMENT:** ***BirthSeq* allows the isolation and investigation of alive cells according to their birthdate, in any kind of tissue and vertebrate species.**

## INTRODUCTION

Embryonic development has been defined as the sequential unfolding of the events of an embryo from which an individual emerges (Barresi and Gilbert, 2000). This process is not homogeneous in time, and most developmental events occur – and need to occur-at specific time points. According to Von Baer’s influential laws of embryology (Abzhanov, 2013), earlier events in embryonic development tend to represent a more general feature of a group of species and usually have a greater influence on the formation of the individual. Development is thus a sequential program that combines the production of increasing numbers of cells with the diversification of different cell types (Duboule, 1994; García-Moreno and Molnár, 2020). Developmental biology is making great progress in identifying both the mechanisms and the role of time in the developmental process.

The impact of developmental time in the formation of the central nervous system is a well-documented example of how a given stem cell progenitor is capable of generating a range of different cell fates along embryonic time (Guillemot et al., 2006; Nieuwenhuys and Puelles, 2016, Klingler et al., 2021; Telley et al., 2016a). The paradigmatic case is that of cortical progenitors in the mammalian telencephalon: one single of these progenitors can generate sequentially a variety of pyramidal neurons for the above lying neocortex, along the cortical formation period (Gao et al., 2014; Telley et al., 2019). Only changing the developmental time when this progenitor divides can modify the cell fate of their daughter cell (García-Moreno and Molnár, 2015). There is evidence of time ruling genomic and proteomic cues. The opening of the chromatin in the Hox cluster requires a specific time (Gaunt, 2015; Montavon and Duboule, 2013). When a Hox gene is activated, it triggers a sequence of genomic interactions that, after a given period of time, enables the expression of the next Hox gene in the cluster. This sequential expression allows the determination of different cell types based on the transcriptional Hox code expressed by the cell (Di Bonito et al., 2013), which is directly related to the time of its formation. As for the proteome, its stability sets a developmental tempo. Proteomic stability encodes for species-specific information about how the spinal cord should develop (Iwata, 2022; Rayon et al., 2020). More recently, time has been shown to play a critical role in establishing a global temporal program for the identity of neuronal subtypes identified in the development of the mouse central nervous system {Formatting Citation}(Sagner et al., 2021). There are probably many more cases in which time is translated into genetic instructions, which in turn drive embryonic development. However, the lack of specific tools allowing the simultaneous exploration of time and gene expression limits our range of exploration.

Here we introduce *BirthSeq*, a solution for the shortage of techniques focused on isolating and analyzing cells based on their birth date. This innovative method enables *in vivo* labeling of cells when these are generated, isolation via FACS at relevant developmental time points, and analysis of cell fate using cutting-edge high-throughput techniques. With a diverse range of administration options, BirthSeq is a versatile tool suitable for any vertebrate species, allowing research on any tissue of the organism. Our tests of the various components of BirthSeq in the brain development of three vertebrate species (mouse, chick, and gecko) have shown evidence of its effectiveness for single-cell RNA sequencing of dated populations of neurons and other organs, as well as novel spatially-resolved transcriptomic techniques. Here we provide first the transcriptomic profile of early-born pallial neurons in mouse; and second, we uncover the genetic identity and anatomical location of early born neurons in the hypothalamus and diencephalon of the chick.

## RESULTS AND DISCUSSION

### FlashTag application in non-mammalian embryos

A permanent labeling system is crucial to reach the goal of studying specifically-dated cell populations. 5-bromo-2’-deoxyuridine (BrdU) and tritiated thymidine, two of the most widely used methods, have a drawback: they make the isolation of live cells impossible, as the staining needed for their visualization is cytotoxic. These two reagents are incorporated into newly synthesized DNA during the S-phase of the cell cycle. However, there is another option that bypasses the DNA synthesis requirement: Carboxyfluorescein succinimidyl ester (CFSE). When incorporated into the cytoplasm of dividing cells, it becomes fluorescent and acts as a birthdating marker. The application of CFSE in the brain ventricles, known as FlashTag (Govindan et al., 2018), has the potential to boost research on birthdated populations, at least for the developing brain. And so, we tested its capabilities in different species.

We replicated FlashTag experiments in chick embryos to test whether it was a useful method in non-mammalian species. With the objective of defining the optimal CFSE concentration to birthdate embryonic chick neuronal populations, we injected different CFSE concentrations in the ventricle of E4 embryos, when pallial neurogenesis begins in chick (Tsai et al., 1981), and analyzed brain cell populations by flow cytometry (FC, **Fig. S1**) 3 days after injection (**Figure S2A**).

Our results showed that neurons birthdated with low CFSE concentrations were almost undetectable and that their brightness was limited (**Figure S2F-I** left panels, **J**). However, this scenario shifted when 5 mM and 10 mM CFSE were used (**Figure S2H,I** left panels). Regarding cell death percentage (**Figure S2F-I** right, **K**), it was maintained below 1% in all tested conditions except for 10 mM CFSE, when it increased to almost 2%.

As 10 mM is the recommended CFSE concentration to label cells in mouse experimental model (Govindan et al., 2018; Telley et al., 2016, 2019), and our FC analysis results showed that this concentration ensures the detection of CFSE+ cells by FC, we decided to check the validity of the pulse-labeling birthdate in chick tissue. The histological analysis revealed that 3 days after the injection, in the VZ of some telencephalic regions like dorsal pallium (DPall) and subpallium (SPall) there were still many CFSE+ neural progenitors (**Figure S2B,C**). Therefore, newborn neurons may become CFSE+ even three days after CFSE administration, making of these injections a less optimal method for high-resolution timed birthdating of neuronal populations in developing chick embryos. Additionally, we observed that CFSE was toxic for chick embryos, as it caused brain hemorrhages and most of the embryos died after the injection (**Figure S2D,L**).

We have revitalized the idea of analyzing cells carrying a DNA mark with our novel method, *BirthSeq*. BrdU has been widely used in the past, but the antigen retrieval process required for detecting it within chromatin kills the cells, making live cell research impossible. To address this issue, BirthSeq is based on 5-etinil-2’-desoxiuridina (EdU), which labels newly generated cells in a similar manner to BrdU, but without the need for harsh treatments on cells to reveal their DNA mark (Salic and Mitchison, 2007). Ideally, EdU can be detected on disaggregated cells while preserving their viability. Our detailed protocol for BirthSeq (**Figure 1A**) involves incorporating EdU into dividing cells through various administration methods, isolating live birthdated cells through FACS sorting, and analyzing them with RNAseq tools.

**Figure 1.**
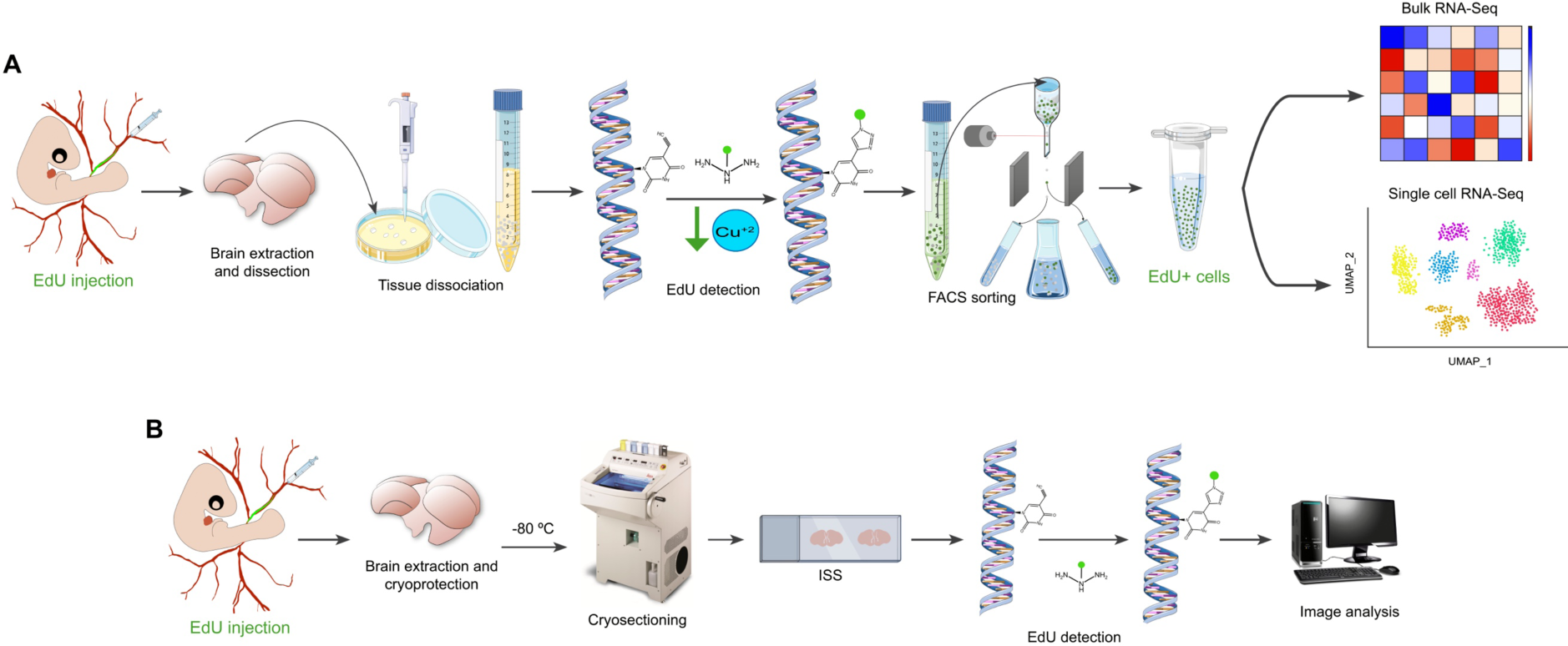
BirthSeq, a novel method to isolate birthdated cells. [A] Schematic representation of *BirthSeq* methodology. BirthSeq consists on the birthdating of any vertebrate tissue by a systemic injection of EdU, followed by the extraction and dissociation of the birthdated tissue of interest. Next, EdU+ cells are detected with a low copper EdU detection reaction mix and isolated by FACS. Finally, the transcriptome of those EdU+ viable cells is analyzed by bulk or single cell RNA sequencing. [B] Schematic representation of *NeurogenesISS,* the modified version of BirthSeq that is compatible with spatial transcriptomics.

In the following sections we tested the suitability and optimal conditions for each of the steps of the BirthSeq protocol.

### *In ovo* administration of EdU labels all dividing and birthdated cells

We conducted a test similar to the one performed on FlashTag to assess the validity of EdU as a reagent for pulse-labeling in birthdating assays. We administered low doses of EdU during the embryonic development of chick, mouse, and gecko embryos and evaluated its birthdating properties. Within 30 minutes of administration, dividing cells in the S-phase region of the ventricular zone of chick embryos acquired the EdU tag (**Fig. 2A**). After 2 hours and 5 hours, the labeled cells moved their nuclei towards the abventricular region of the ventricular zone where they eventually divided during the M-phase (**Fig. 2B,C**). Importantly, 5 hours after administration, there was no new incorporation of EdU in the S-phase region, indicating that EdU at these low doses is rapidly washed from the ventricular zone and is an effective pulse-labeling factor in chick embryos. The administration of low doses of EdU did not result in an increase in embryo mortality (n>100) and did not generate the aberrant isochronic clusters that have been reported with high-dose BrdU in chick embryos (Rowell and Ragsdale, 2012).

**Figure 2.**
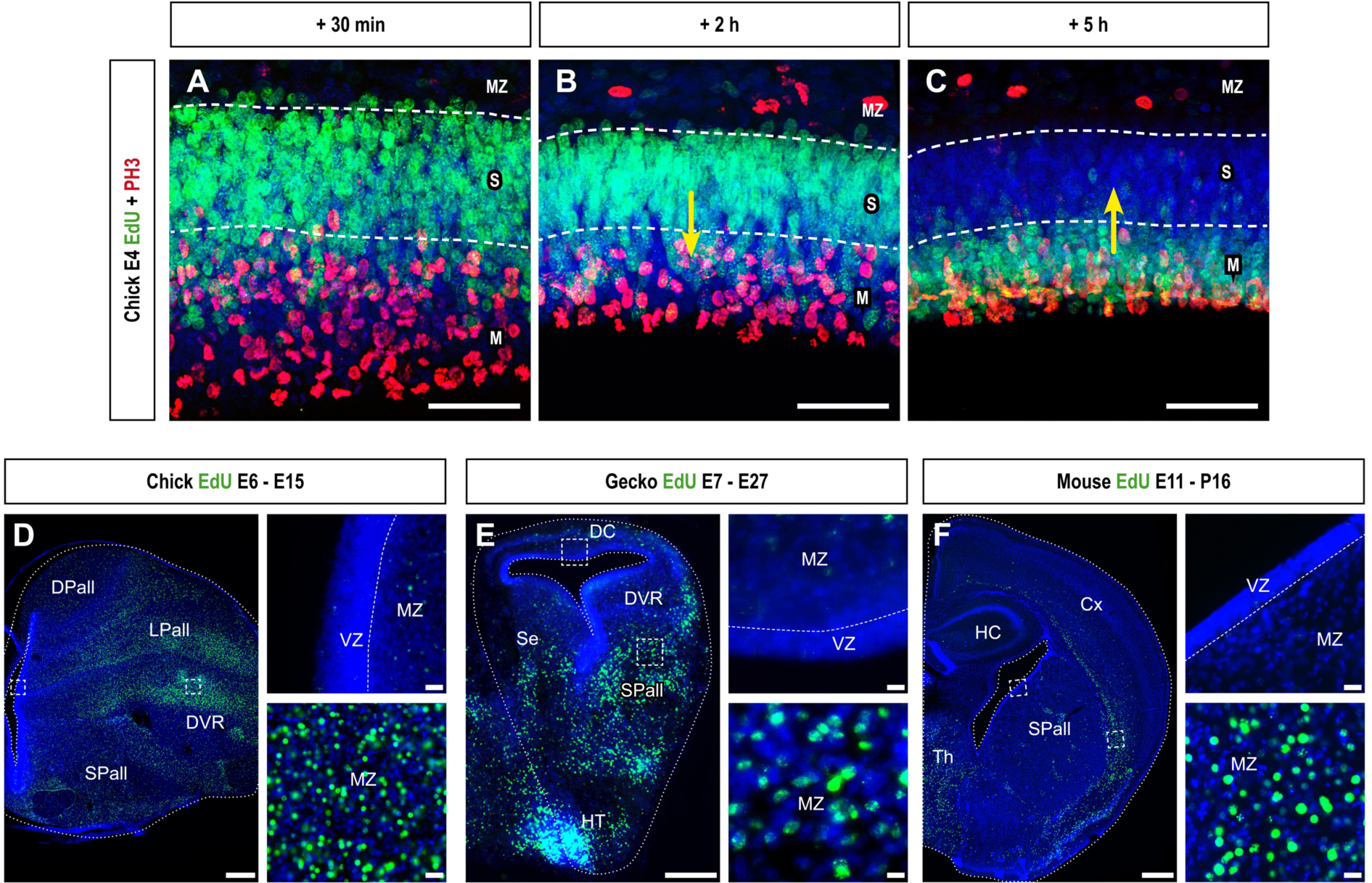
*In vivo* EdU administration is an effective birthdating pulse-labeling assay. [A-C] Short-term birthdating of chick ventral pallial cells was performed at E4. [A] Thirty minutes after birthdating, EdU+ cells appeared at the most basal region of the neuroepithelium. [B] Two hours after the EdU injection, part of the EdU+ cells had started their migration to the ventricular surface, following their interkinetic nuclear movement. [C] Finally, five hours after EdU administration, many of the EdU-labeled cells were undergoing mitosis (counterstained with PH3) in the most apical region of the neuroepithelium. In addition, the S-phase region appeared mostly blank of EdU labelling, as no new EdU was being incorporated at this time after the injection. [D-F] Long-term examples of EdU incorporation and labeling in brain coronal sections of E15 chick, injected at E6 [D]; E27 gecko, injected at E7 [E]; and P16 mouse, injected at E11 [F]. In all of the researched species there is an evident lack of EdU-labeled cells in the VZ of the telencephalon and the presence pf many scattered EdU+ cells throughout the MZ. *For nomenclature, refer to the list of abbreviations. DAPI counterstain in blue. Dashed white lines in [A-C] demarcate the neuroepithelial region where neural stem cells duplicate their DNA (S-phase, S); whereas in [D-F] demarcate anatomical boundaries. Yellow arrows in [B] and [C] mark the cellular movement. Scale bars, 50 µm [A-C], 500 µm [D, F], 100 µm [E], 25 µm [insets of D, F], and 5 µm [insets of E]*.

We tested the long-term viability of EdU as a birthdating reagent in mouse, chick, and gecko embryos, and our results showed that the birthdating was successful in all three species (**Fig. 2D-F**). Our examination of their brains several days after the administration of EdU revealed a differential staining pattern that was consistent with brain development. The neurons generated during the time of administration were labeled, while many other cells were not. This confirmed that the pulse-labeling was effective, as there was no remaining EdU mark in the progenitor cells days after the injection [**Fig. 2D-F**, (Rueda-Alaña and García-Moreno, 2022)].

To maximize the potential of *BirthSeq*, EdU can be administered via different methods to the embryo. In mouse experiments, it was administered through intraperitoneal injections via the pregnant dam. Chick and gecko embryos were mostly administered with intracardiac or intravenous injections. These three administration vias proved to be advantageous as they are systemic, meaning that the EdU molecule reaches all cells in the organism. It allows all organs to be researched with a single systemic dose, providing a more reliable birthdating analysis. In the case of the brain, cells divide on several regions and neuroepithelial strata, some are not available for a labeling from the ventricle. We sorted out this limitation as systemic EdU also reaches dividing cells in subventricular positions, making the birthdating even more comprehensive. Intraventricular injections, although not systemic, were also able to reach all dividing cells in the developing brain.

Our results demonstrate that EdU is a highly effective birthdating reagent for histological analysis, capable of being used in all amniote species and for the study of any developing organ or tissue.

### Labeling and FACS isolation of viable EdU birthdated cells

Next, we evaluated the feasibility of using EdU labeling for the FAC-Sorting of living cells. Our initial efforts to isolate live birthdated cells were unsuccessful, primarily due to the toxicity of the labeling reaction mixture to reveal the EdU labeling. Since the reagents used in the labeling reaction are well-known, we made modifications to the labeling protocol to maintain cell viability while still achieving successful fluorescent labeling. We experimented with different exposure times to the reaction mixture and reaction temperatures (**Figure S3**), but the effective improvement was reducing the amount of Cu (II) in the reaction mixture (**Figure 3**).

**Figure 3.**
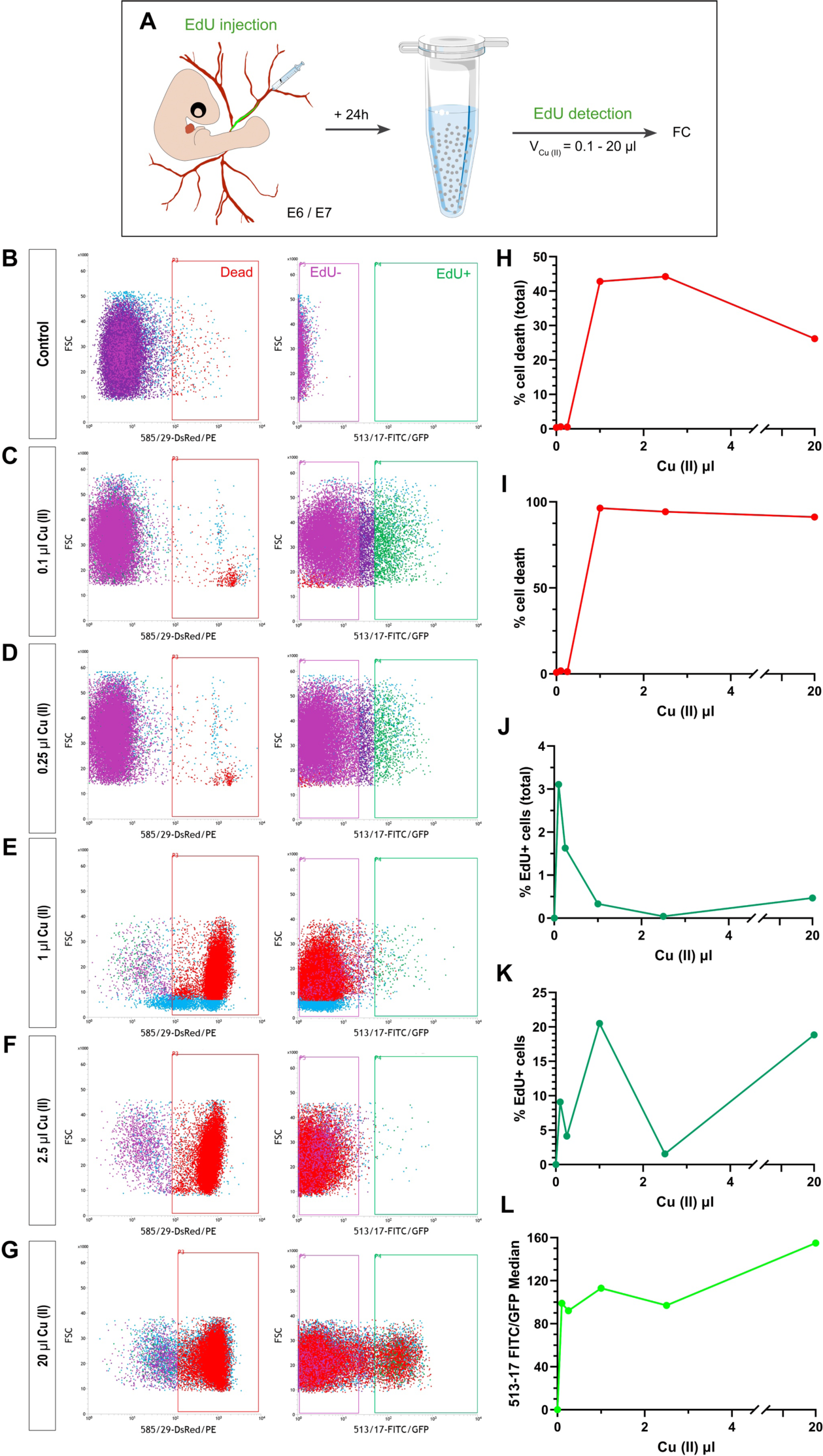
The reduction of Cu (II) concentration in the EdU detection cocktail increases cell viability without losing EdU detection capacity. [A] Experimental design of the FC analysis of dated chick embryonic neurons after Cu (II) volume variations in the EdU detection cocktail. [B-G] FC profiles of dissociated chick embryonic neural cells 1 day after EdU injection and EdU detection with 0 µl [B], 0.1 µl [C], 0.25 µl [D], 1 µl [E], 2.5 µl [F], and 20 µl [G] of Cu (II). [H-L] Graphic representation of FC analysis. Relative proportion of dead [H-I] and EdU+ [J-K] cells of total and parental populations were analyzed. EdU+ cell brightness was assessed by 513-17 FITC/Median [L]. FC analysis demonstrated that Cu is toxic for the cells, as shown by the high cell death rates observed in the samples revealed with high Cu (II) concentrations and its dramatic reduction after lowering the concentration. However, reducing copper concentration did not diminish the EdU detection capacity of the cocktail, making it possible to isolate EdU+ viable cells. *Dead cells (P3, red) were defined as PI^+^ in FSC versus dsRed dotplot (left panel) while EdU+ (P4, green) and EdU-(P5, purple) cells were identified as GFP^+^ and GFP^-^ in FSC versus GFP dotplot (right panel). Data represented in the graphs was obtained from a single cytometry experiment, and analyzed samples contained 3 pooled brains*.

EdU “click” technology relies on the creation of a covalent bond. This chemical reaction is catalyzed by Cu (I), which is created in situ from Cu (II) obtained from CuSO4 (Rostovtsev et al., 2002; Salic and Mitchison, 2007; Tornøe et al., 2002). It is well known that Cu (II) is toxic for the cells, as it reduces mRNA transcription and promotes RNA degradation (Halliwell and Gutteridget, 1984; Halliwell et al., 1992). However, there is evidence that, in fixed cells, an 85% reduction in Cu (II) concentration was able to replicate the standard EdU detection capacity (Ng et al., 2017). Hence, we decided to analyze by FC chick brain birthdated samples after modifying Cu (II) concentration in the EdU cocktail (**Figure 3**).

Cell death percentages decreased proportionally with Cu (II) reduction, until reaching almost undetectable levels after 80-fold and 200-fold Cu (II) reductions (**Figure 3B-G** left, **H-I**). At the same time, we observed that EdU+ cells were still distinguishable although their brightness was slightly reduced (**Figure 3B-G** left, **L**). Further analysis showed that samples revealed with low Cu (II) were enriched with EdU+ cells when compared to the rest of the samples (**Figure 3B-G** left, **J**), being these an expected consequence of the cytotoxicity reduction. In addition, we observed that the EdU parental population (EdU+ viable cells with the size and shape of interest) was smaller in those samples revealed with low copper (**Figure 3B-G** left, **K**), which, together with brightness loss, could be indicating that the Cu (II) concentration decrease is reducing the capacity to detect EdU.

Considering all these results, we decided to work with a 200-fold Cu (II) concentration reduction (as in **Fig. 3C**). In spite of the fact that with this Cu (II) concentration our EdU detection capacity was reduced, its cytotoxicity was irrelevant. We would still be able to isolate birthdated cells that have divided only once after the injection (so the EdU of their DNA would not have been diluted and would be enough to be detected), maintaining a high survival rate. For our particular experimental setting, this is actually an advantage, allowing a more exact birthdating.

As mentioned above, Cu (II) can interfere with RNA expression (Halliwell and Gutteridget, 1984; Halliwell et al., 1992; Ng et al., 2017) Consequently, we decided to examine if the new EdU cocktail (containing low Cu (II) concentration) was altering cell gene expression. For that, we purified by FACS a non-revealed control population and a treated population, and compared their gene marker expression by RT-qPCR **(Fig. S4**). First, FACS results showed that the treatment was not modifying population size and that it was safe for cell viability (**Fig. S4B-E**). Second, RT-qPCR analysis showed that the mRNA expression was not altered in the treated sample (**Fig. S4F**) so the new EdU cocktail did not seem to modify the transcriptional machinery of the cells. This RT-qPCR analysis was sufficient for our purpose, and let us continue in our optimization of BirthSeq for scRNAseq and other *omics* technologies.

Once the EdU detection protocol was optimized, the next step was to verify if it was sensitive enough to selectively isolate birthdated cells. In order to do that, we compared the mRNA expression of two neural cells types that, in the moment of the isolation, were at two different stages of maturation in the E7 chick brain: neural progenitors (birthdated 30 minutes before isolation) and immature neurons (birthdated 3 days before isolation) (**Figure 4A-E**). We found that, even though there is only statistical significance in Sox2 expression difference, neural progenitors tended to have a higher mRNA expression of progenitor markers than immature neurons (**Figure 4F**). This difference of expression tendency shifted its orientation when neural markers were analyzed (**Figure 4G**). Taken together, these results suggested that the protocol was valid to selectively isolate EdU+ cells.

**Figure 4.**
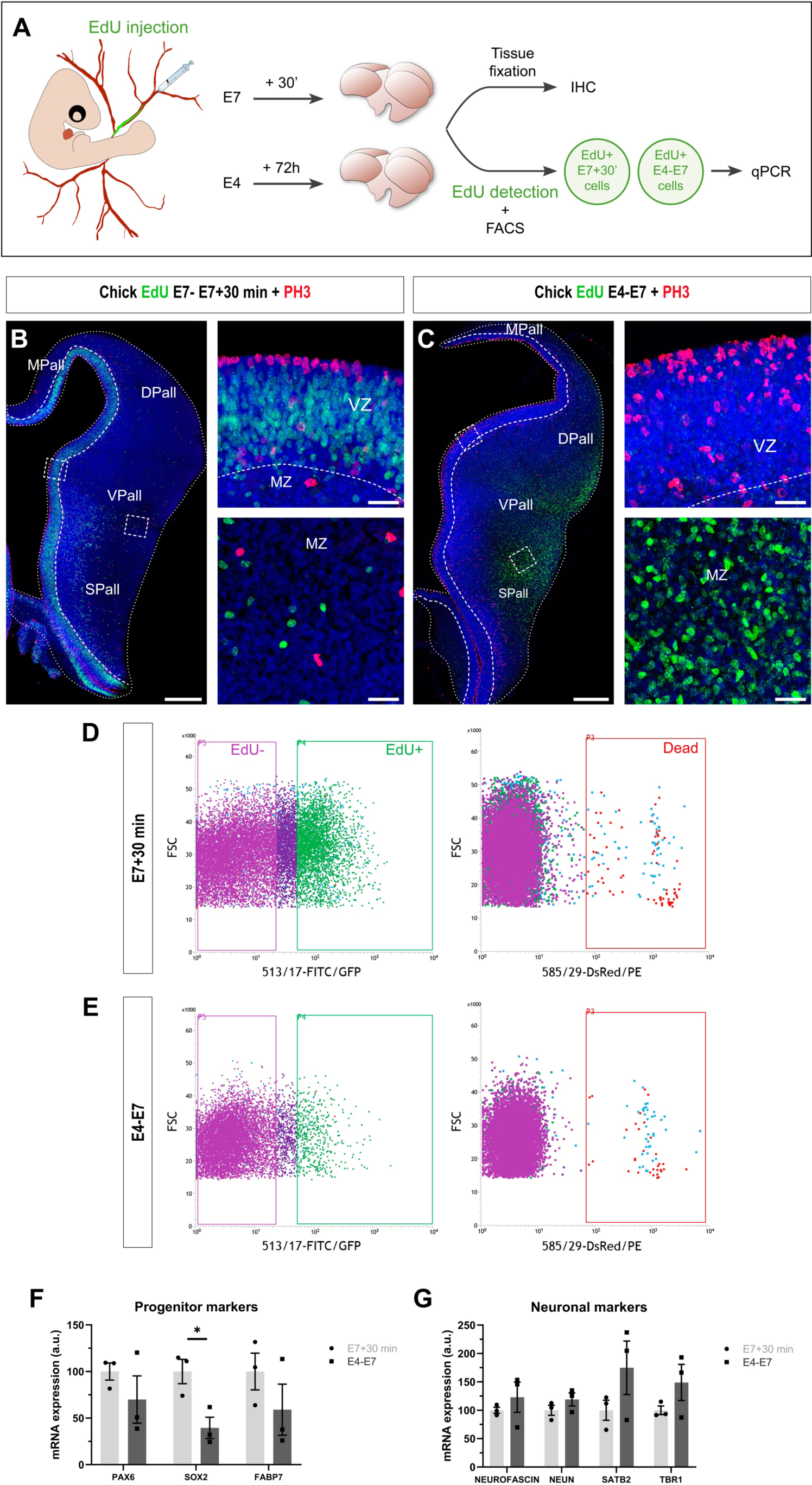
EdU-mediated cell isolation is specific for birthdated cells. [A] Experimental design used to analyze gene expression of birthdated neuronal progenitors and immature neurons. [B-C] Coronal sections and high-power views of E7 chick telencephala injected with EdU 30 minutes [B] or 3 days [D] before the analysis. Pictures clearly show that E7 chick embryos birthdated 30 minutes before the analysis only have EdU+ progenitors located in the VZ, while there is a lack of EdU-labeled cells in the MZ. However, three days after being birthdated, EdU+ cells present in E7 chick telencephalon are located in the MZ and correspond to immature neurons. [D-E] FC profiles of birthdated chick neuronal progenitors [D] and neurons [E]. [F-G] Expression of progenitor [F] and neuronal [G] markers in neuronal progenitors (E7 + 30 min) vs immature neurons (E4-E7) cells by RT-qPCR. Progenitor cells expressed higher levels of progenitor markers, whereas neuronal markers were more expressed in immature neurons than in neural progenitors. RPL7 was selected as a reference gene. *For nomenclature, refer to the list of abbreviations. DAPI counterstain in blue. Dashed white lines demarcate anatomical boundaries. Scale bars, 250 µm [B-C], and 25 µm [B-C insets]. EdU+ (P4, green) and EdU-(P5, purple) cells were identified as GFP+ and GFP-in FSC versus GFP dotplot (left panel), whereas dead cells (P3, red) were defined as PI+ in FSC versus dsRed dotplot (right panel). *p<0.05. Student’s t test in [D]. Bars represent mean ± SEM. Dots show individual data. Each sample contained 4 pooled brains*.

Finally, we wanted to ratify the versatility of the method and verify that the new EdU cocktail was valid to detect various other types of birthdated cells. Consequently, EdU was intravenously injected in chick embryos to reach systemically all cells of the embryo (**Fig. S5**). Three days after the injection, embryonic hearts and limbs were dissociated and EdU was detected on those two tissue samples (**Fig. S5A**). FC analysis of parental populations showed that cell death percentage was low in both cardiac and limb cells (**Fig. S5B-C** left**, D**), suggesting that EdU cocktail is safe for cells obtained from different tissues. As well, we were able to visualize (and, therefore, isolate if needed) bright EdU+ populations in both samples (**Fig. S5B-C** right**, E-F**), which, once again, validated the new EdU cocktail to detect birthdated cells.

### BirthSeq cells can be used for bulk or single-cell RNA sequencing

One of the final goals of BirthSeq is to conduct high-throughput sequencing experiments on cell populations of which birthdate is known. We tested the applicability of BirthSeq to single cell RNA sequencing experiments. We utilized the protocol for extracting E12.5-generated cortical cells from P3 mouse pups (**Figure 5A,B**). At this stage of postnatal development, the cortex already harbors populations of neurons and glial cells (**Figure 5B**). This is relevant because it is known that neurons are more sensitive to the pressure and aggressive conditions of FACS sorting than glial cells (Pan and Wan, 2020; Ryan et al., 2021). When we applied BirthSeq to the brains of these mice, we obtained the expected outcome in the single cell RNA dataset: several populations of glial cells were present in the dataset as they survived better under the BirthSeq conditions, as well as the expected and birthdated populations of neurons (**Figure 5C-D**). Specifically, the injection of EdU at E12.5 labeled Cajal-Retzius cells, immature deep layer cortical neurons, and other pallial populations known to be generated at this time point, including GABAergic interneurons (**Figure 5E,F**). Additionally, we found earliest-generated layer IV neurons, which may correspond to the latest divisions of EdU+ progenitors (**Figure 5E,F**). All these cells exhibited the typical markers of the populations they belonged to (**Figure 5F**), thereby confirming that BirthSeq was capable of generating a single cell RNA sequencing dataset enriched in the populations of cells generated shortly after the EdU injection. Cajal-Retzius cells are the best example since they comprise the minor population of cortical neurons and are known to disappear from the tissue at postnatal stages (Meyer, 1999). Any cortical dataset from P3 mouse cortex would reveal very few of these cells (Di Bella et al., 2021); whereas, in our birthdated dataset, it is a highly enriched cluster (**Figure 4G**). This data confirms BirthSeq as a novel method to isolate and analyze birthdated populations using single cell RNA sequencing methodologies. This is one of the first examples where single cell RNA sequencing is applied to cells for which birthdate is known a priori, and the first method that is not limited to a specific cell type, organ, or species.

**Figure 5.**
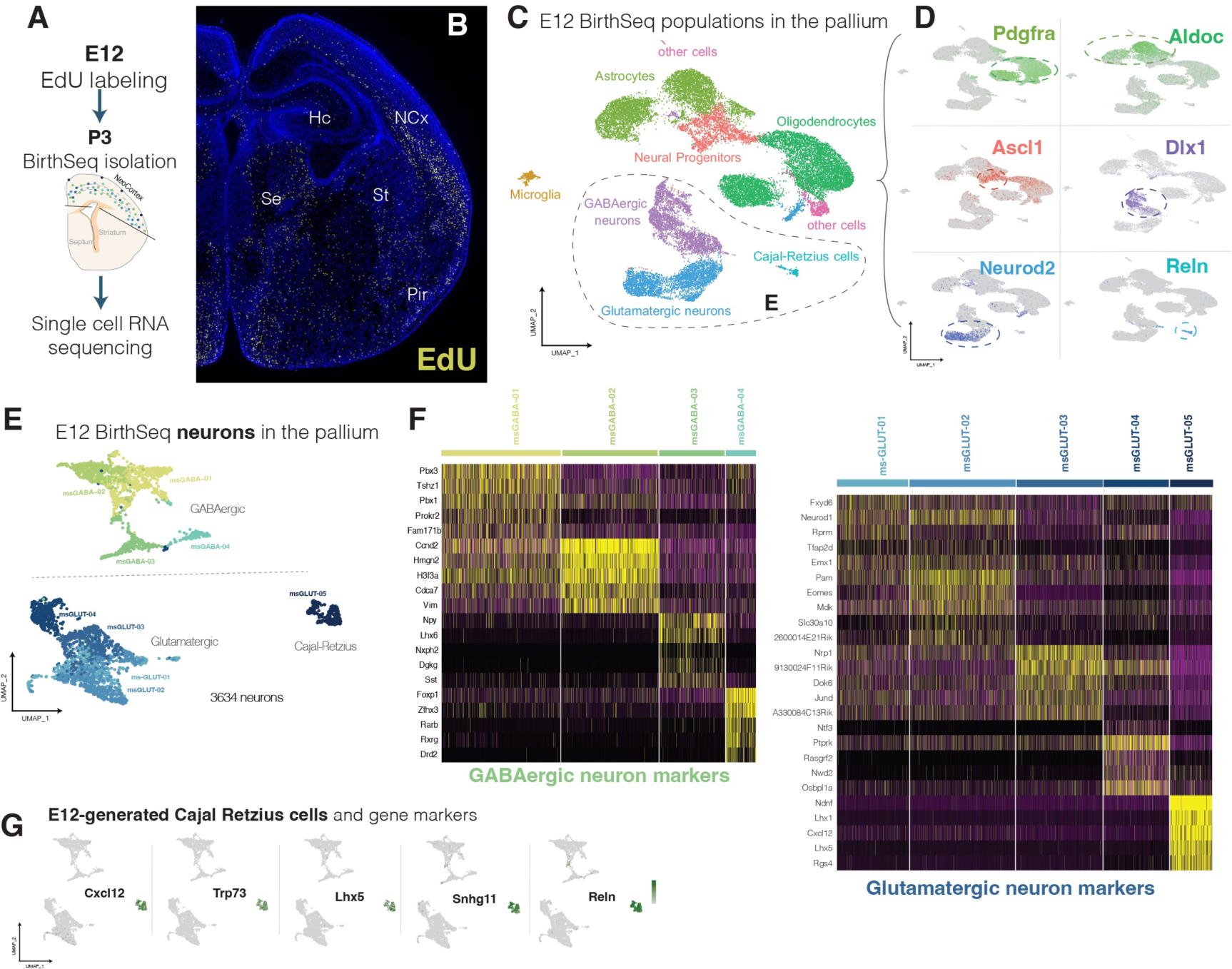
BirthSeq allows single cell RNA sequencing of birthdated populations in the mouse brain. [A] Experimental design. Pregnant dams were injected at E12, when neurogenesis is starting in the neocortex of mouse embryos. After surgery, embryos were allowed to grow up to P3, when we applied the BirthSeq protocol followed by scRNAseq. [B] EdU staining (yellow) of P3 mouse telencephalon showing the location of birthdated cells within the tissue in a littermate; DAPI counterstain is shown in blue. [C] UMAP distribution of E12 birthdated cells after BirthSeq. All main neural populations were identified. The insets at the right panel shows representative gene plots for marker genes of each main cell type. [E] UMAP distribution of the neuronal populations only, including glutamatergic and GABAergic neurons. [F] Heatmap of gene markers for each of the 9 neuronal clusters identified and generated at E12, organized in 4 GABAergic and 5 glutamatergic clusters. [G] Feature plots of different Cajal-Retzius cell markers to show the enrichment of this early-born population in the sequenced dataset.

### *NeurogenesISS*: BirthSeq along spatially-resolved transcriptomics

EdU labeling can be detected by fluorescent microscopy, which allowed us to combine BirthSeq with spatial-omics techniques. To test the compatibility of the EdU label with in situ sequencing (ISS)(Ke et al., 2013)(Lee at al., 2023), we worked on an E15 chick brain sample treated with BirthSeq at E4 and used ISS to detect the expression of 84 genes at single-cell resolution (**Figure 6**). At the end of the ISS detection cycles, we revealed the EdU label using click-chemistry. Given our focus on the generation of diencephalic neurons, we named this combined tool *NeurogenesISS*. Our use of both techniques proved successful, enabling us to determine the transcriptomic profile and brain location of the diencephalic and hypothalamic neurons generated at E4 (**Figure 6A**).

**Figure 6.**
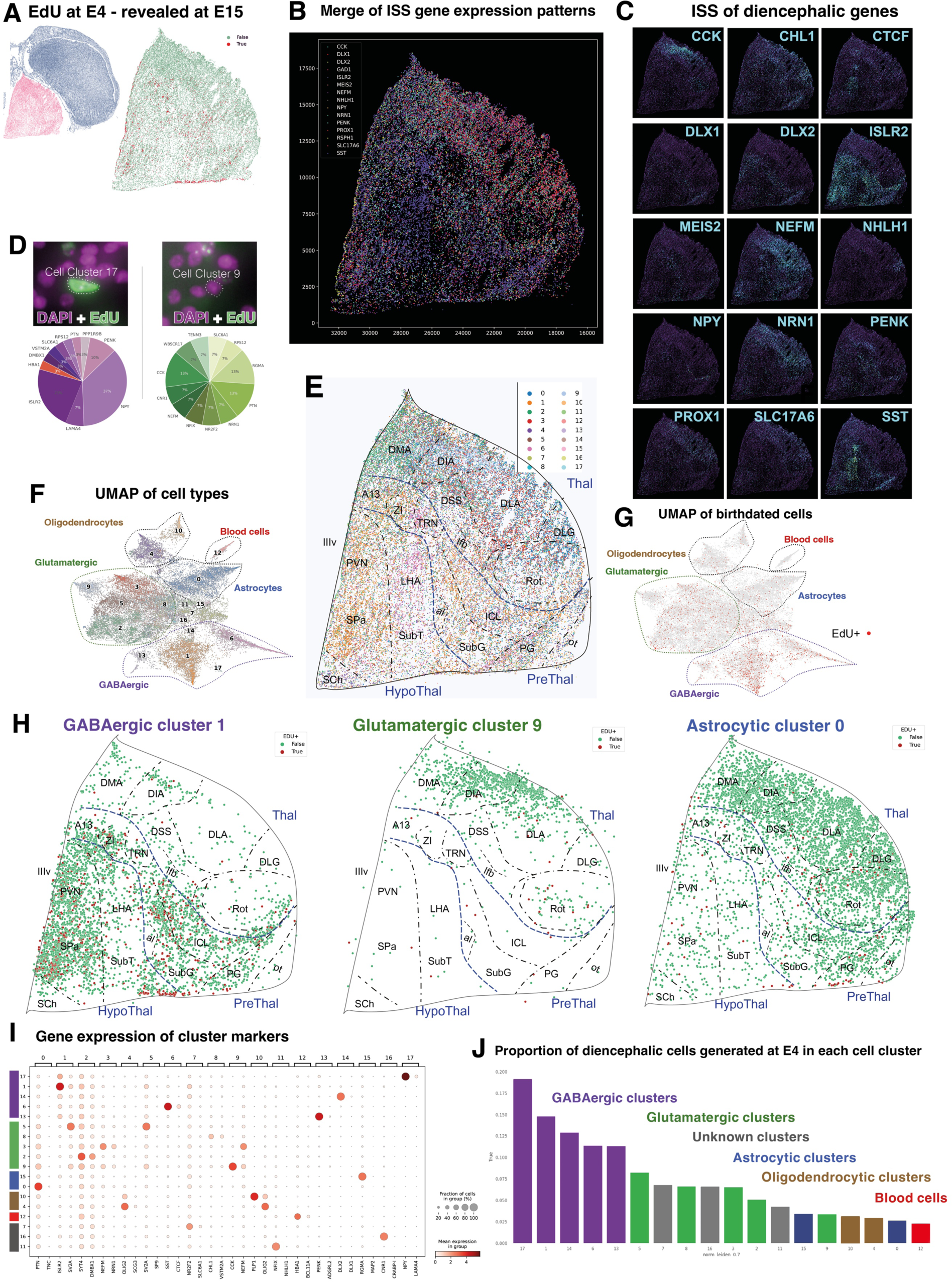
*NeurogenesISS* resolves spatial transcriptomics of birthdated populations in the chick brain. [A] Anatomical location of the brain region under research, the diencephalon (highlighted in pink), in a coronal section. The embryos were injected at E4, and the EdU staining was revealed at E15, depicted by the red dots in the right panel (with the diencephalon isolated). [B] Merge image of 16 genes from the panel obtained through ISS (in situ sequencing) reveals broad regional differences between the thalamus, prethalamus, and hypothalamus. [C] Example of the gene expression pattern of 16 relevant genes detected after in situ sequencing. [D] A high magnification detail image illustrates the ISS mapped reads of two types of cells: One EdU(+) cell from cluster 17, and EdU(-) cell from cluster 9. [E] Distribution of 18 cell clusters in the diencephalon detected after clustering of the cells, informed by the quantitative expression of 84 analyzed genes. It shows the identification of diencephalic and hypothalamic nuclei, guided by ISS gene expression and the distribution of cell clusters. [F] UMAP distribution of cells belonging to the 18 clusters, grouped by major cell type. [G] Equivalent UMAP distribution, highlighting in red the cells that were EdU+ (generated at E4). [H] Distribution of 3 cell classes in the diencephalic section, organized by major cell type. Green dots represent cells of a given cluster, while red dots represent cells of the cluster that were EdU+ (generated at E4). [I] Dot plot representing the expression of ISS gene markers on each cell cluster. [J] Proportion of cells within each cluster that were generated at E4, following the administration of EdU.

We initiated our analysis by determining the expression pattern of the 84 genes tested within the tissue sample (**Figure 6B,C**). We then segmented the tissue into individual cells based on the nuclear signal, and assigned to each cell a set of neighbouring ISS signals situated within its segmentation mask (**Figure 6D,E**). At this stage the data consist of a cell by gene expression matrix, which can be mainly treated in the same way as a single-cell RNA dataset. We proceeded to perform clustering of the diencephalon cells based on their gene expression similarity (**Figure 6D-G**). Our analysis produced a comprehensive spatial map of cell types, detailing the location of each cell type within the diencephalon (**Figure 6F**). In total, we identified 18 clusters corresponding to 34,717 total mapped cells (**Figures 6D,F,G and S6**). Thalamic clusters, with their glutamatergic nature, were easily identifiable, as were GABAergic neurons of the prethalamus and hypothalamus. These cells belong to major classes of cells such as: 5 classes of GABAergic neurons mainly distributed across the prethalamus and regions of the hypothalamus (*ISLR2, NPY, DLX2, GAD1, SST, PENK*), 5 classes of glutamatergic neurons mainly located within the limits of the thalamus and regions of the hypothalamus (*NEFM, CCK, SYT4, SV2A, CHL1, SLC17A6*), 2 types of astrocyte cells with a different distribution pattern (*PTN, RGMA, AQP4*), and another 2 types of oligodendrocytes of varied maturation degree (*OLIG2, PLP1, PDGFRA*). Additionally, one cluster belonged to blood cells (*HBA1*), and 3 clusters of unknown identity were recognised.

Finally, by means of the EdU administered at E4 we determined the diencephalic and hypothalamic neurons generated at that specific timepoint. We extracted the EdU intensity for each cell, and defined as EdU-positive all the cells that had a median intensity superior to a given conservative threshold. We then extracted the identity and location of all the EdU-positive cells. This enabled us to visualize the location and genetic profile of chick neurons generated at E4 (**Figure 6H,I**). Our findings revealed that GABAergic cells of the prethalamus and hypothalamus were the most abundant population of cells generated at the E4 stage (**Figure 6J**). A good example is the distribution of cluster 1, comprising GABAergic cells located in the prethalamus and hypothalamus. In this cluster identified by enriched expression of *ISLR2*, up to 15% cells were generated at E4 as labeled by EdU (**Figure 6J**). These populations were specifically located in the reticular thalamic nucleus, the pregeniculate nucleus, the intercalate nucleus of the prethalamus, and the paraventricular and subparaventricular nuclei of the hypothalamus. Also, nearly 20% of all *NPY+* GABAergic cells, located in the surroundings of the rotundus nucleus (cluster 17 in **Figure S6**), were generated at E4. On the other hand, the percentage of glutamatergic neurons generated at E4 was consistently lower than that of GABAergic cells, ranging from 2.5% in the case of *CCK+* neurons of the dorsal anterior nucleus of the thalamus to 7.5% of other glutamatergic thalamic neurons labeled with *SV2A* (clusters 9 and 5, respectively, in **Figure S6**). As expected from an early labeling timepoint, the percentage of glial cells generated was the lowest among the neural cells detected, with a maximum of 3% of either astrocytes or oligodendrocytes of the four classes detected being generated as early as E4 (**Figure 6G,K**). This data represents the first instance of coupling spatially-resolved transcriptomics and birthdating analysis.

The groundbreaking potential of BirthSeq and its spatial-omics variant NeurogenesISS has yielded highly compelling results, establishing their position as powerful tools to explore the critical role of developmental timing in generating diverse cell populations. With their versatility, these techniques can be effectively applied across various vertebrate species and cell types from any organ, paving the way for further research in the dynamic fields of proteomics and epigenomics. BirthSeq tools are versatile and can be used in conjunction with various scRNAseq platforms. They can also be combined with either scATACseq (Baek and Lee, 2020) or multiomics approaches to uncover changes in the epigenetic profiles of cells during differentiation. When combined with CITE-seq (Stoeckius et al., 2017) BirthSeq tools enable the identification of cell populations and their ages based on their surface protein expression. Additionally, when used alongside Perturb-seq (Dixit et al., 2016), BirthSeq tools can help pinpoint specific genes that play a crucial role in the formation of these dated cell populations. BirthSeq development represents a significant breakthrough in our understanding of the intricate mechanisms underlying cellular differentiation and developmental processes, promising to unlock new avenues of scientific discovery.

## MATERIALS AND METHODS

### EXPERIMENTAL ANIMALS

All animal experiments were approved by the University of the Basque Country (UPV/EHU) Ethics Committee (Leioa, Spain) and the Diputación Foral de Bizkaia, and conducted in accordance with personal and project licenses in compliance with the current normative standards of the European Union (Directive 2010/63/EU) and the Spanish Government (Royal Decrees 1201/2005 and 53/2013, Law 32/107). Fertilized chick eggs (*Gallus gallus*) were purchased from Granja Santa Isabel (Córdoba, Spain). They were incubated at 37.5 °C in humidified atmosphere until required developmental stage. The day when eggs were incubated was considered embryonic day (E)0. Adult C57BL/6 mice (*Mus musculus*) were obtained from a mouse breeding colony at Achucarro Basque Center for Neuroscience (Spain). Animals were housed in a 12/12 h light/dark cycle (8 AM, lights on), at constant temperature (19-22 °C) and humidity (40-50%); and provided with ad libitum food and water. The day when the vaginal plug was detected was referred to as E0. Fertilized ground Madagascar gecko eggs (*Paroedura picta*) were obtained from a breeding colony at Achucarro Basque Center for Neuroscience (Spain). Adult geckos were housed in a 12/12 h light/dark cycle (8 AM, lights on, 27 °C/ 8 PM light off, 22 °C) and provided with ad libitum food (live crickets) and water. Gecko eggs were incubated at 28 °C in a low humidified atmosphere until required developmental stage. The day when eggs were harvested from the terrarium was considered E0.

### IN OVO PROCEDURES AND BIRTHDATING EXPERIMENTS

Manipulation of chick and gecko embryos was performed as previously described (Rueda-Alaña and García-Moreno, 2022; Rueda-Alaña et al., 2018). Briefly, eggs were incubated in a vertical position at either 37.5 or 28 °C. Birthdating molecules were administered via intraventricular, intracardiac or intravenous injections for chick embryos or via intraventricular injection for gecko embryos, by mouth-pipetting using a fine pulled-glass needle. Drops of sterilized Hank’s balanced salt solution (HBSS, HyClone Cytiva) were added to prevent embryonic dehydration, eggs were sealed and embryos were incubated until needed.

#### EdU birthdating

EdU was diluted in sterile phosphate buffered saline (0.1 M PBS, pH 7.6) with 0.1% Fast Green FCF (Sigma), and the dose was adjusted for each species and developmental stages. Chick embryos were injected with a single dose of 1 µl EdU: E4 embryos were injected with 2.5 µg/µl EdU; E6 and E7 embryos were injected with 5 µg/µl EdU. Gecko E7 embryos were administered a single 0.5 µl dose of 25 ng/µl EdU. Pregnant mice dams were intraperitoneally injected with 150 mg/kg EdU at E11 and E12.

#### CFSE birthdating

CFSE (CellTrace™ CFSE, Life Technologies) was diluted in DMSO (Sigma-Aldrich) with 0.1% Fast Green FCF. For the FlashTag FC experiments, E4 chick embryos were intraventricularly injected with 1 µl of 1-10 mM CFSE. For histology and tissue dissociation optimization experiments, E4 chick embryos were injected intraventricularly with 1 µl of 10 mM CFSE.

### BRAIN TISSUE PROCESSING: PERFUSION, FIXATION AND SECTIONING

#### For immunohistochemistry

All the brains were fixed in 4% paraformaldehyde (PFA, Sigma-Aldrich) diluted in PBS. Gecko brains and the chick embryonic brains up to E7 were collected in ice-cold HBSS and fixed by immersion in PFA for 24 h, transferred to PBS and kept at 4 °C. P3 postnatal mouse pups, and E15 chick embryos were anesthetized by hypothermia after immersion of either pup or the chick egg in ice. P16 postnatal mouse pups were deeply anesthetized with 2.5% 2,2,2-tribromoethanol (Avertin; Sigma-Aldrich). Then they were transcardiacally perfused with PBS followed by PFA. Brains were removed and postfixed, with the same fixative, for 3 h at RT, then transferred to PBS and kept at 4 °C.

Brains were serially sectioned in the coronal plane using a Leica VT1200S vibrating blade microtome (Leica Microsystems GmbH, Wetzlar, Germany) and stored at 4 °C until use. All mouse and gecko brains were cut in 50 µm-thick sections. Embryonic chick brains were cut in 60 µm-thick sections, to prevent tissue breakdown.

#### For in situ sequencing

E15 chick embryos were anesthetized by hypothermia. Brains were extracted in ice-cold HBSS, cryopreserved in 30% sucrose, fresh frozen in optimal cutting temperature media (OCT, Tissue-Tek®) and stored at -80 °C until sectioning. Tissue was sectioned in the coronal plane, under RNAse-free conditions, at 12-16 µm thickness with a Leica CM1950 cryostat (Leica Microsystems GmbH, Wetzlar, Germany) and collected on SuperFrost Plus microscope slides (VWR). Slides were stored at -80 °C until processing.

### IMMUNOHISTOCHEMISTRY AND BIRTHDATING STAINING

Single immunohistochemical reactions were performed as described previously (Rueda-Alaña and García-Moreno, 2022; Rueda-Alaña et al., 2018). Briefly, sections were incubated with blocking and permeabilization solution (PBS containing 0.5 % Triton-100X (Sigma-Aldrich) and 3 % bovine serum albumin (BSA, Sigma-Aldrich)) for 3 h at RT and then incubated overnight with the primary antibodies (diluted in the same solution) at 4 °C and agitation. After the incubation, the primary antibodies were removed. Sections were washed five times with 0.5 % PBT (0.5% Triton-100X in PBS) for 10 min, and once with the blocking and permeabilization solution for 10 min. Next, sections were incubated with fluorochrome-conjugated secondary antibodies diluted in the same solution for 2 h at RT in the dark. Finally, sections were washed with 0.5 % PBT twice for 10 min.

In birthdated animals, at the end of the immunohistochemical process, EdU molecule was detected in the sections according to the manufacturer’s protocol using “click-it” chemistry (EdU Cell Proliferation Kit, baseclick GmbH, Neuried, Germany). Finally, sections were washed with 0.5 % PBT and mounted on SuperFlost slides (Epredia™) with Mowiol mounting medium (Mowiol® 4-88, Sigma-Aldrich).

All sections were counterstained with DAPI (1:1,000; Sigma-Aldrich). The following antibodies and EdU azides were used: rabbit anti-PH3 (1:1,000; GeneTex), Alexa Fluor 488 Goat anti-rabbit (1:1,000; Invitrogen), 6-FAM azide (1:500; Lumiprobe) and Sulfo-Cyanine 5 Azide (1:500; Lumiprobe).

#### Image capture and analysis

Images were collected using a Leica SP8 laser scanning confocal microscope (Leica, Wetzlar, Germany) or a 3DHistech Panoramic Midi II digital slidescanner (3DHistech, Hungary).

Images obtained with the confocal microscope were acquired with the LAS X software. The signal from fluorophores was collected sequentially-green light and far-red light were collected together. Same image parameters (laser power, gain and wavelengths) were maintained for images from each slide and adjusted for new animals. For big brain sections, tile-scan images were composed. All images shown are projections from z-stacks ranging from 10 to 35 µm of thickness, typically 20 µm. Images obtained with the slidescanner microscope were acquired in single layer mode, using the 20X objective and collecting the signal from fluorophores sequentially.

### BIRTHSEQ OPTIMIZATION PROCEDURES

#### Tissue dissociation and sample preparation

All chick samples for this set of experiments were dissociated using an adaptation of previously described protocols (Beccari et al., 2018; Huettner and Baughman, 1986; Moussaud and Draheim, 2010). Briefly, brains were extracted, collected in ice-cold HBSS and pooled. Tissue was cut into small pieces and placed into full chick enzymatic solution (see details in **table 1**). Samples were incubated at 37 °C for 15-20 min, manually homogenized by carefully pipetting and filtered through a 40 µm nylon strainer (Fisher) to a 15 ml Falcon tube quenched by 4 ml of 25% Fetal Bovine Serum (FBS, Gibco) in HBSS. Afterwards, cell suspensions were centrifuged at 200g for 10 min at RT.

**Table 1.** Summary of buffer, solution and cocktail recipes.

| Buffer / Solution | Composition |
| --- | --- |
| **STOCK enzymatic solution** | 116 mM NaCl (Sigma-Aldrich), 5.4 mM KCl (Sigma-Aldrich), 26 mM NaHCO<sub>3</sub> (Sigma-Aldrich), 1 mM NaH<sub>2</sub>PO<sub>4</sub> (Sigma-Aldrich), 1.5 mM CaCl<sub>2</sub> (Sigma-Aldrich), 1 mM MgSO<sub>4</sub> (Sigma-Aldrich), 0.5 mM EDTA (Invitrogen), 25 mM glucose (Sigma-Aldrich), 1 mM L-cysteine (Sigma-Aldrich). |
| **Full chick enzymatic solution** | STOCK enzymatic solution containing 150 U/ml DNase I (Invitrogen) and 50 $\mu\text{g/ml}$ proteinase K (Sigma-Aldrich). |
| **Full mouse enzymatic solution** | STOCK enzymatic solution containing 150 U/ml DNase I (Invitrogen) and 11.3 U/ml papain (Sigma-Aldrich). |
| **Sorting buffer** | 25mM HEPES (Sigma-Aldrich), 5mM EDTA, 1% BSA, in HBSS |
| **Traditional EdU detection reaction cocktail** | 1:10 Reaction buffer (10X), 1:25 Catalyst solution (CuSO<sub>4</sub>), 1:500 Dye azide (10 mM), 1:10 Buffer additive (10X) |
| **New EdU detection reaction cocktail** | 1:10 Reaction buffer (10X), 1:5,000 Catalyst solution (CuSO<sub>4</sub>), 1:500 Dye azide (10 mM), 1:10 Buffer additive (10X) |

#### FC and FACS

FC was performed on a FACS Jazz (BD Biosciences). Cells were scanned for propidium iodide (PI, (BD Pharringen); excitation, YG561 laser; detection, BP585/29 filter) and 488 signal (excitation, B488 laser; detection, BP513/17 filter). Same voltage parameters were maintained in all the samples of an experiment, and adjusted for each experiment.

FACS was performed using a FACS Jazz, with a 100 µm nozzle. Cells were scanned for PI (excitation, YG561 laser; detection, BP585/29 filter) and EdU-488 signal (excitation, B488 laser; detection, BP513/17 filter) to collect PI-/EdU+ and PI-/EdU-cell populations. Sorting pressure was always 27 PSI.

#### Definition of cell populations and hierarchies

This is the definition of the several **total** cell populations (P) that were gated in the samples (**Figure S1**):

- The <u>P2 population</u> comprises all cells of an adequate size (defined by high FSC) and complexity (defined by low SSC) in the sample (**Figure S1A**).
- The <u>P6 population</u> comprises all singlets (individualized cells) present in the sample (**Figure S1B**).
- The <u>P3 population</u> comprises all dead cells (positive for PI at 585/29-DsRed/PE) present in the sample (**Figure S1C**).

`o` The <u>NOTP3 population</u> was formed by all the live cells (negative for PI at 585/29-DsRed/PE) present in the sample. This population was defined by a Boolean gating of P3 (**Figure S1C**).
- The <u>P4 population</u> comprises all CFSE+ or EdU+ green cells (positive at 513/17-FITC/GFP) present in the sample (**Figure S1D**).
- The <u>P5 population</u> comprises all CFSE- or EdU-cells (negative at 513/17-FITC/GFP) present in the sample (**Figure S1D**).

In order to analyze and isolate specific cell types, parental populations were defined by establishing a strict cell population hierarchy during FC and FACS procedures:

1. First, P2 population was selected, excluding cell debris and nuclei.
2. Second, P6 parental population was defined by selecting the singlets among the P2 population.
3. Third, P3 parental population was defined by identifying the dead cells among the P6 parental population.
4. Fourth, NOTP3 parental population was defined by excluding the P3 parental population from the analysis.
5. Fifth, P4 parental population was defined by selecting the green fluorescent cells among the NOTP3 parental population.
6. Sixth, P5 parental population was defined by selecting the non-green fluorescent cells among the NOTP3 parental population.

All the hierarchy and parental populations are detailed in **Figure S1E-F**.

#### CFSE-mediated isolation of birthdated chick embryonic neurons

For the FlashTag method experiments (Govindan et al., 2018; Telley et al., 2016b; Telley et al., 2019), E4 chick embryos were intraventricularly injected with 1-10 mM CFSE. Three days later, brains were extracted and dissociated. Obtained cell pellet was resuspended in 500 µl of sorting buffer (see details in **table 1**), filtered through a 50 µm cell strainer and analyzed by FC . Analyzed samples are detailed in **table 2**.

**Table 2.**
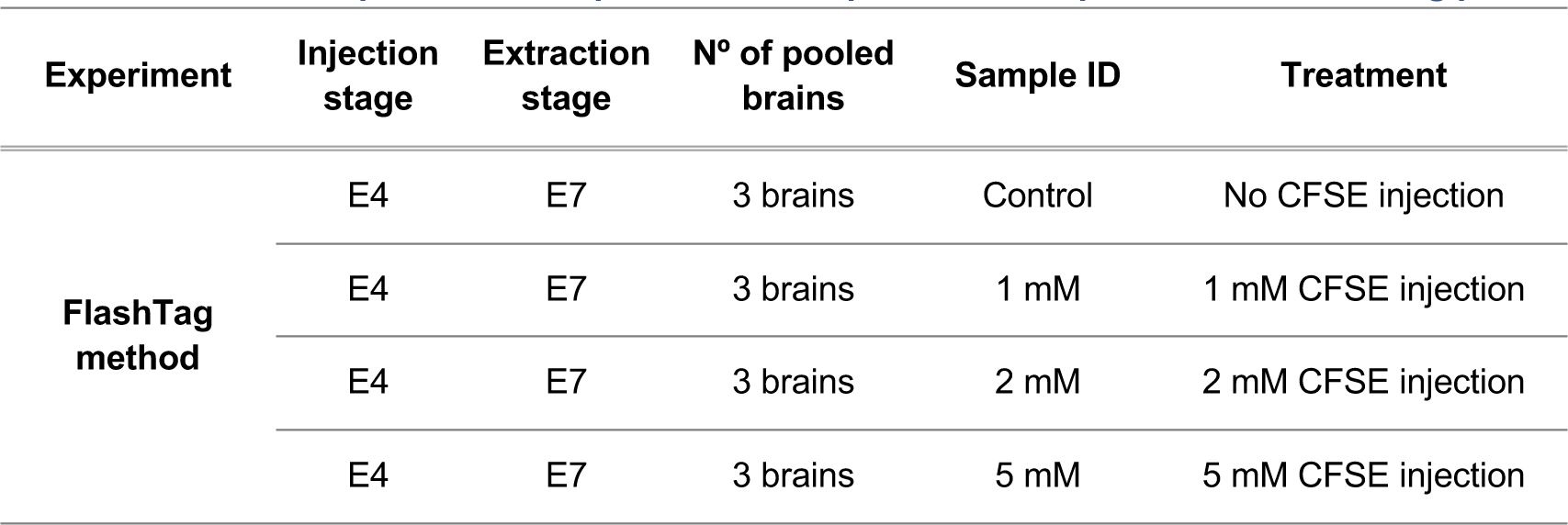
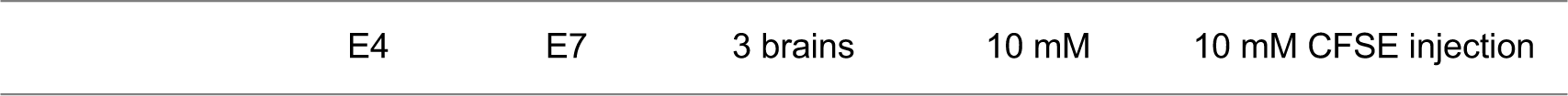
Detailed description of the experimental samples used to optimize the birthdating protocol.

#### Optimization of time, temperature and reagent concentration for EdU detection

E7 chick embryos were intravenously injected with EdU and one day later (at E8) brains were extracted and dissociated. After dissociation, different protocols were used for EdU detection (see details in **table 3**). After EdU detection, cells were washed by centrifugation at 200g for 10 min and resuspended in 500 µl of HBSS. The washing step was repeated twice. Finally, cell pellets were resuspended in 500 µl of sorting buffer, filtered through a 50 µm strainer and analyzed by FC.

**Table 3.** Detailed description of the experimental samples used to optimize the EdU detection conditions.

| Injection stage | Extraction stage | N° of pooled brains | Sample ID | EdU detection cocktail composition | EdU incubation conditions |
| --- | --- | --- | --- | --- | --- |
| E7 | E8 | 3 brains | Control | HBSS | 30 min at RT in the dark |
| E7 | E8 | 3 brains | Treatment A | Traditional EdU detection reaction cocktail | 30 min at RT in the dark |
| E7 | E8 | 3 brains | Treatment B | Traditional EdU detection reaction cocktail | 10 min at RT in the dark |
| E7 | E8 | 3 brains | Treatment C | Traditional EdU detection reaction cocktail | 30 min at 15 °C in the dark |
| E7 | E8 | 3 brains | Treatment D | Modified traditional EdU detection cocktail with reagent concentration reduced to half | 30 min at RT in the dark |
| E7 | E8 | 3 brains | Treatment E | Modified traditional EdU detection cocktail with reagent concentration reduced to half | 30 min at 15 °C in the dark |

#### Optimization of the copper concentration in the EdU detection reaction cocktail

The optimal CuSO4 · 5H2O concentration for EdU detection was identified on an experiment performed on E7 and E8 chick embryonic brains that were intravenously injected with EdU 24 h before. Birthdated brains were extracted and dissociated. After dissociation, pelleted cells were resuspended in EdU detection reaction cocktail; which contained variable copper volumes (see details in **table 4**). After EdU detection, samples were washed twice with 500 µl of HBSS and resuspended in 500 µl of sorting buffer. Finally, samples were analyzed by FC after being filtered through a 50 µm strainer.

**Table 4.** Detailed description of the experimental simples used to optimize the catalyst (Cu^+2^) concentration for EdU detection.

| Injection stage | Extraction stage | N° of pooled brains | Sample ID | EdU detection cocktail composition | EdU incubation conditions |
| --- | --- | --- | --- | --- | --- |
| E7 | E8 | 3 brains | Control | HBSS | 30 min at RT in the dark |
| E6 | E7 | 3 brains | 0.1 μl Cu (II) | EdU detection cocktail with a 200-fold Cu (II) reduction | 30 min at RT in the dark |
| E6 | E7 | 3 brains | 0.25 μl Cu (II) | EdU detection cocktail with an 80-fold Cu (II) reduction | 30 min at RT in the dark |
| E7 | E8 | 3 brains | 1 μl Cu (II) | EdU detection cocktail with a 20-fold Cu (II) reduction | 30 min at RT in the dark |
| E7 | E8 | 3 brains | 2.5 μl Cu (II) | EdU detection cocktail with an 8-fold Cu (II) reduction | 30 min at RT in the dark |
| E7 | E8 | 3 brains | 20 μl Cu (II) | Traditional EdU detection reaction cocktail | 30 min at RT in the dark |

#### Validation of the EdU detection reaction cocktail

With the goal of checking if the EdU detection protocol was interfering with gene expression, E4 chick embryos were intravenously injected with EdU. Two days later, brains were extracted, dissociated and incubated in the new EdU detection cocktail at RT for 30 min. Next, cells were washed thrice by centrifugation at 200g for 10 min and resuspended with HBSS. Finally, samples were passed on a 50 µm cell strainer, and all the viable PI negative singlets with the size and shape of interest, regardless of being EdU+ or EdU (P1 parental population), were isolated by FACS. P1 population was collected in Buffer NR (NZYTech) containing 1% β-mercaptoethanol (Sigma-Aldrich) and stored at -80 °C until processing. Analyzed samples are detailed in **table 5**.

**Table 5.** Detailed description of the experimental samples used to validate Birth-Seq.

| Experiment | Injection stage | Extraction stage | N° of pooled brains | Sample ID | EdU detection cocktail composition | Isolated cell type |
| --- | --- | --- | --- | --- | --- | --- |
| Validation of the EdU detection cocktail | E4 | E6 | 3 brains | - EdU reaction cocktail | HBSS (no detection) | Viable singlets with correct size and shape |
|  | E4 | E6 | 3 brains | + EdU reaction cocktail | New EdU detection cocktail | Viable singlets with correct size and shape |
| Isolation of different types of birthdated neural cells | E7 | E7+30 min | 4 brains | E7 + 30 min | New EdU detection cocktail | Neural progenitors |
|  | E4 | E7 | 4 brains | E4 – E7 | New EdU detection cocktail | Immature neurons |
| Isolation of birthdated cells from different non-neural tissues | E4 | E7 | 2 hearts | Heart | New EdU detection cocktail | Birthdated cardiac cells |
|  | E4 | E7 | 4 limbs | Limb | New EdU detection cocktail | Birthdated limb cells |

#### Isolation of birthdated neural cells

Both immature neurons (birthdated three days prior isolation) and neuronal progenitors (birthdated 30 min before the extraction) were isolated. Briefly, E7 chick brains were extracted and dissociated (see details in Tissue dissociation). Next, EdU was detected by incubating the samples in the new EdU detection cocktail for 30 min at RT. Samples were then washed thrice by centrifugation at 200g for 10 min and resuspended in HBSS. Finally, clogs were removed by filtering the sample through a 50 µm strainer and EdU+ viable cells were FAC-Sorted. Isolated cells were collected in Buffer NR containing 1% β-mercaptoethanol and stored at -80 °C until processing. Analyzed samples are detailed in **table 5**.

#### Isolation of birthdated cardiac and limb cells

Birthdated cardiac and limb cells were obtained from E7 chick embryos that were intravenously injected with EdU at E4. Tissue was collected on ice-cold HBSS, dissociated and samples were incubated in the new EdU detection cocktail for 30 min at RT. Next, samples were washed three times by centrifugation at 200g for 10 min, resuspended in HBSS and filtered. Finally, samples were analyzed by FC. Analyzed samples are detailed in **table 5**.

### RNA ISOLATION, RETROTRANSCRIPTION AND QPCR

#### FAC-Sorted cell RNA isolation and retrotranscription

RNA from FAC-Sorted EdU+ cells (neural cells >650,000) was isolated by NZY Total RNA Isolation kit (NZYTech) according to the manufacturer’s instructions. RNA was quantified in QubitTM 4 Fluorometer (Invitrogen) with the Qubit® RNA HS Assay Kit (Invitrogen).

RNA was retrotranscribed using the NZY First-Strand cDNA Synthesis Kit (NZYTech) following the manufacturer’s instructions in Veriti Thermal Cycler (Applied Biosystems).

#### RT-qPCR

RT-qPCR was performed following MIQE guidelines (Bustin et al., 2009) on a CFX96 Touch Real-Time PCR Detection System (Bio-Rad). Three replicates of 1.5 µl of each cDNA were amplified using NZYSpeedy qPCR Green MasterMix (2x, NZYTech). The amplification protocol was 3 min at 95°C for denaturing; and 45 cycles of 10 s at 95°C and 30 s at 60°C for annealing. The expression level ratios of target genes to housekeeping genes were calculated, and these ratios were normalized with the expression levels of the non-treated control group or with the expression levels of the progenitor group.

#### Primers

Primers (Sigma-Aldrich) were designed to amplify exon–exon junctions using PrimerBlast (NIH) to avoid amplification of contaminating genomic DNA, and their specificity was assessed using melting curves. Amplification efficiency was calculated for each pair of primers using the software LinRegPCR (Ramakers et al., 2003; Ruijter et al., 2009). Primer sequences are listed in **table 6**.

**Table 6.**
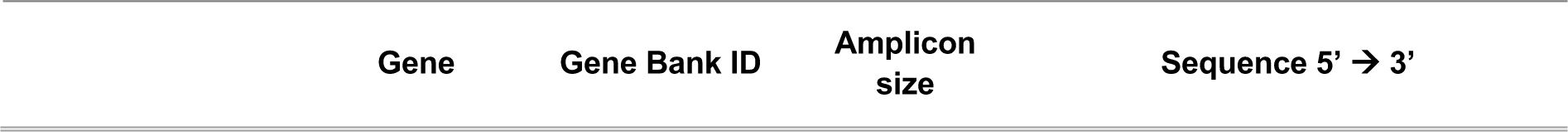
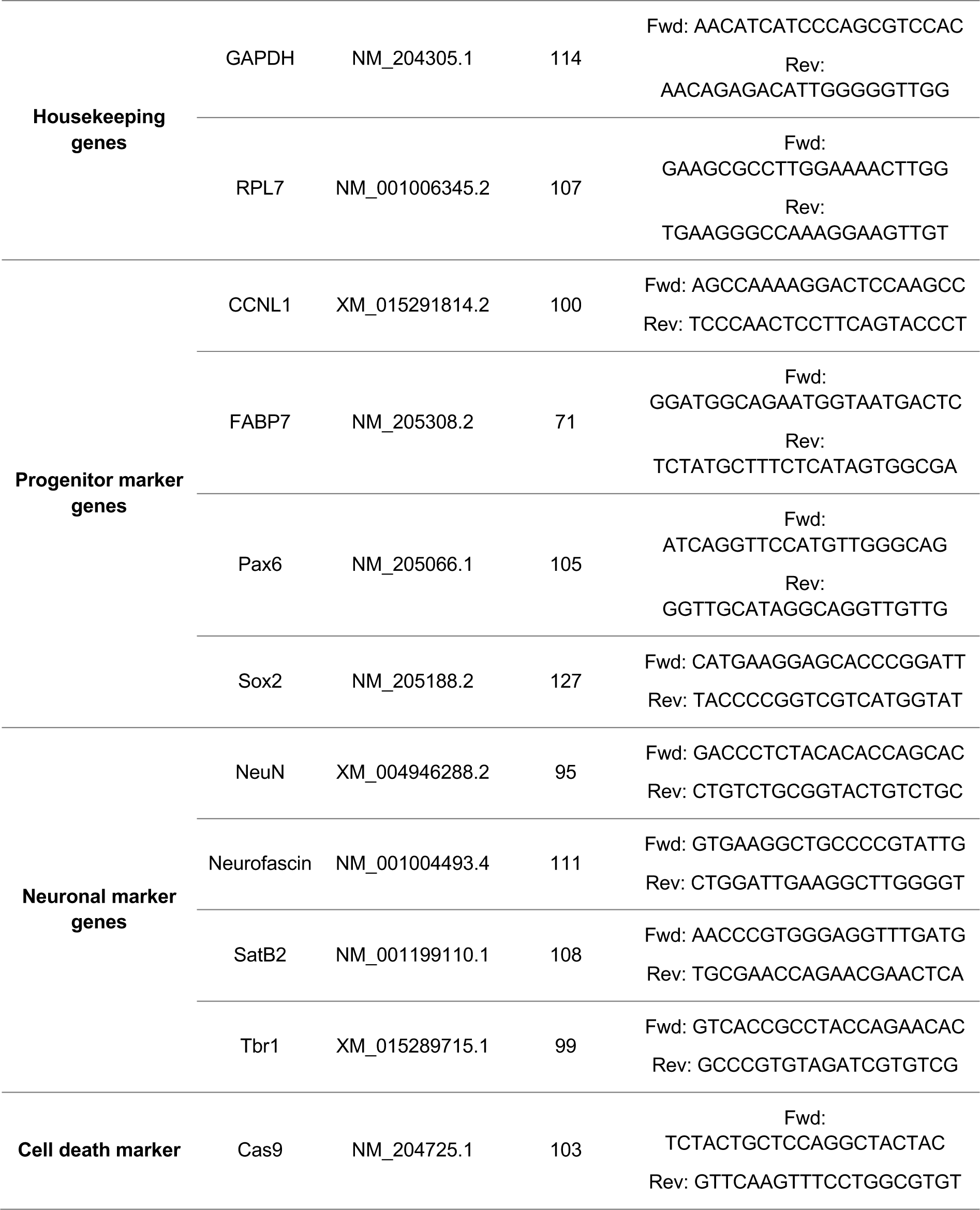
List of used primers.

Two independent reference genes were compared (*GAPDH* and *RPL7*) and their expression remained constant independently of time and treatments (data not shown), validating their use as reference genes. In all experiments, the pattern of mRNA expression was similar using the assigned couple of reference genes, and in each experiment the reference gene that rendered lower intragroup variability was used for statistical analysis.

### SINGLE CELL RNA SEQUENCING (scRNAseq)

#### Cell preparation

We applied *BirthSeq* for the isolation of birthdated cells on P3 mice injected with EdU at E12.5. Briefly, at P3 brains were extracted in ice-cold HBSS, the pallial region was microdissected, meninges were removed under a stereomicroscope, collected in ice-cold HBSS, pooled and cut into small pieces. Next, tissue was incubated in full mouse enzymatic solution (see **table 1**) for 15 min at 37 °C. Following enzymatic digestion, tissue was manually homogenized by carefully pipetting. Then, tissue clogs were removed by filtering the cell suspension through a 40 µm nylon strainer to a 15 ml Falcon tube containing 4 ml of 25% FBS in HBSS. Dissociated cells were then centrifuged at 200g for 10 min at RT, pellets were resuspended in 500 µl of the new EdU detection reaction cocktail (see **table 1**) and incubated at RT for 30 min in the dark. Following reaction, cells were washed thrice by centrifugation at 200g for 10 min and suspended in 1 ml of HBSS then passed on a 50 µm nylon strainer. PI-/EdU+ cells, gated to include only the top 10% brightest cells were finally analyzed by FC and FAC-sorted.

FAC-sorted cells were centrifugated at 200g for 10 min and resuspended in HBSS, at a concentration of 1200 cells per µl. Cell concentration and viability was verified using a TC20 Automated Cell Counter (BioRad). Next, cells were processed for single cell GEM formation following the standard Chromium Single Cell 3’ v3.1. Briefly, cells were placed at the Chromium chip with the beads and reagents for the RNA capture and cDNA amplification. After obtaining the emulsion, cDNAs were amplified, purified and QCed (visualized and quantified by Bioanalyzer 2100). Post-amplified cDNAs, labelled with individual cell barcodes, were fragmented, ligated to sequencing adapters and amplified with dual indexes. After that, the PCR product was purified, quantified by Qubit 2.0 and the profiles visualized in a Bioanalyzer 2100.

The 10X libraries were sequenced in a HiSeq 4000 and a NovaSeq 6000 for an approximated 50.000 reads per cell. When aiming to identify subtle difference among closely related cell types it is important to perform deep sequencing of the libraries.

#### Single cell pre-processing

10X *CellRanger* v6.0.2 was employed for alignment and demultiplexing of *FASTQ* files to obtain feature-barcode matrices. The genome used as reference was mm10-2020-A. Afterwards, data matrices were imported to *R* (v4.1.0), where *Seurat* (v4.1.0) (Hao et al., 2021) was employed to further analysis, as describe in their vignettes (https://satijalab.org/seurat/).

The cell-cycle phase was determined by *CellCycleScore*() function, using “Rrm2”, “Pcna”, “Slbp”, “Wdr76”, “Mcm5” as S-phase genes and “Cenpf”, “Tpx2”, “Hmgb2”, “Ube2C”, “Bub1B”, “Top2A”, “Cenpe”, “Tacc3”, “Bub1”, “Aurka”, “Cdc20” as G2M genes, and the mitochondrial percentage of each cell was calculated with *PercentageFeatureSet*(*pattern= “MT-”).* A manual identification of poor-quality clusters was performed and those cells were excluded for next steps. Samples of equal temporal stages were merged into one Seurat object were subject to normalize and scale, and regress-out cell-cycle variation.

#### Cluster identification

To group cells by transcriptome similarity, expression data was linearly reduced into principal components (*RunPCA*, default parameters). A shared nearest neighbor (SNN) graph was calculated from the first principal components that explains almost 90% of the variability or the %variability explained by the next PC is less than 5% with *FindNeighbors*, default parameters. Based on this graph, the Louvain algorithm allows to detect communities or clusters with multi-level tuning of the resolution parameter (*FindClusters*, resolution (0.05, 0.1, 0.2, 0.3, 0.4, 0.5, 0.6, 0.8, 1.0, 1.2, 1.8, 2.4)). This resolution varied from lower values, to identify general cell types (e.g. “glutamatergic neurons”); to higher values, to identify cell subtypes (e.g. “MGE-derived GABAergic interneurons”). Finally, cells were represented into a two-dimensional space by the non-linear dimensional reduction technique *Uniform Manifold Approximation and Projection* (*RunUMAP*) and also *t-Distributed Stochastic Neighbor Embedding (tSNE)*.

To identify the neurobiological cell identity of each cluster, differential expression analysis was carried out among clusters (*FindAllMarkers*, *min.pct = 0.25, logfc.threshold = 0.25*). Scientific literature, *in situ* hybrization databases (Allen Brain Institute, Mouse Genome Informatics), single cell equivalent experiments from the literature (Bandler et al., 2022; Di Bella et al., 2021; Li et al., 2020; Telley et al., 2019) were used to assign cell type identities. As said before, there was a first identification at low resolution, where neurons and neural progenitors were subsetted as cells of interest; and a high resolution, where only cells of interest were classified into more specific subtypes. The genes used for the first and second assignments are displayed in several heatmaps and *FeaturePlots* (**Fig. 5**).

### IN SITU SEQUENCING

#### ISS Library preparation

Slides with tissue sections were thawed and brought to room temperature for 5 minutes, then washed twice with PBS (room temperature) and progressively dehydrated with a 70% ethanol bath for 2 minutes, followed by a 100% ethanol bath for 2 minutes, and finally air dried. Secure-Seal hybridization chambers (Grace Biolabs, #621502) were attached to the microscope slides, to cover the samples, and filled with PBS-Tween 0.5%, followed by a PBS wash. The sections were fixed in fresh 3% PFA in PBS for 5 minutes, followed by 3 PBS washes. The tissue was permeabilized for 5 minutes in 0.1N HCl and washed twice with PBS.

To better discriminate cell types in the brain section, the selection of genes to be assessed by ISS was led by a previous scRNAseq experiment. We then chose significant genes, due to their specific and high expression, for the differentiation of the several cell types. A probe solution was prepared according to the following recipe: 2x SSC, 10% Formamide and 10 nm of each padlock probe. The sequence for all the probes can be found in the Supplementary Table 1. The solution was incubated on the samples overnight at 37 C. The next day we washed the unhybridized excess probes by 2 washes of 10% formamide in 2x SSC, followed by 2 washes in 2x SSC. After removing the last SSC wash, a ligation mix was prepared as follows: 50 mM Tris-HCl (pH 7.5), 1 mM DTT, 100 µM ATP, 2 mM MgCl2, 50 nM of RCA primer, 1U/µL of RiboProtect (Blirt, #RT35) and 0.5 U/ µL of T4 RNA Ligase 2 (NEB, # M0239L). The ligation mix was introduced to the SecureSeal chamber and incubated on the samples for 2 hours at 37 degrees Celsius. After ligation, the samples were washed twice with PBS and an amplification mix was prepared as follows: 50 mM Tris-HCl (pH 8.3), 10 mM MgCl2, 10 mM (NH4)2SO4, 5% Glycerol, 0.25 mM dNTPs, 0.2 µg/mL of BSA and 1 U/µL of Phi29 polymerase (Blirt, #EN020). The amplification reaction was carried out overnight at 30 C. The next day, the samples were washed 3 times with PBS and L-probes (or bridge probes) were incubated at 10 nM each for 30 minutes in 2xSSC and 20% Formamide. Excess probes were washed out with 2 washes in 2x SSC and detection oligos and DAPI were incubated for 30 minutes in the same conditions as L-probes. Excess detection oligos were washed out with 2 washes in 2x SSC. TrueBlack (Biotium #23007) was applied to quench background fluorescence, according to the manufacturer’s instructions. Samples were mounted in SlowFade Gold (ThermoFisher, S46936), and cyclical imaging was performed. After each cycle of imaging, Lprobes and detection oligos were stripped with two washes of 3 minutes in 100% formamide, followed by 5 washes in 2x SSC. Hybridisation of L probes and detection oligos for the following detection cycle was performed as above.

#### ISS Image acquisition

Imaging was performed using a standard epifluorescence microscope (Leica DMI6000) connected to an external LED source (Lumencor® SPECTRA X light engine). Light engine was set up with filter paddles (395/25, 438/29, 470/24, 555/28, 635/22, 730/40). Images were obtained with a sCMOS camera (2048 × 2048, 16-bit, Leica DFC90000GTC-VSC10726), automatic multi-slide stage, and Leica Apochromat 40× (HC PL APO 40x/1.10 WATER, 11506342) objective. The microscope was equipped with filter cubes for 5 dyes separation (AF750, Cy5, Cy3, AF488 and DAPI) and an external filter wheel (DFT51011).

Each region of interest (ROI), corresponding to a hemisphere of each one of the sections, was marked and saved in the Leica LASX software, for repeated imaging. Each ROI was automatically subdivided into tiles, and for each tile a z-stack with an interval of 0.5 micron was acquired in all the channels. The tiles are defined to have a 10% overlap at the edges. The images were saved as thousands of individual tiff files with associated metadata.

#### EdU staining

At the end of the last ISS detection cycle, we stripped all the L-probes and detection oligos by 2 washes in 100% formamide, followed by 5 washes of 2x SSC. We then proceeded to the click-labeling of EdU with an Alexa Fluor 488 dye, using the Click-iT EdU Cell Proliferation Kit for Imaging (ThermoFisher #C10337), following the manufacturer’s instructions. Imaging was done as above, and the images were processed as if they were a normal ISS cycle.

#### Image data processing

The raw images and the associated metadata from the microscopes were fed into the pre-processing module of a custom analysis pipeline (Lee et al, 2023). The module transforms the images into a format suitable for decoding, executing the following steps: first the images are maximum Z-projected. The resulting 2D projected images (”tiles”) are simultaneously stitched and registered across imaging cycles, using the ASHLAR (Muhlich et al., 2022) software. ASHLAR captures the metadata and places the tiles correctly in the XY space before starting the alignment step. During the process, the 10% overlap is also removed to produce stitched images. Finally, the aligned stitched images are sliced again into smaller tiles, to allow a faster and computationally efficient decoding. The resliced aligned tiles were denoised using Content-Aware Image Restoration (CARE) (Weigert et al., 2018), applying a custom denoising model, trained in the lab specifically on ISS data. The resulting images were then converted into the SpaceTX format(Axelrod et al., 2021), and fed into the Starfish Python library for decoding of image-based spatial transcriptomics experiments (https://github.com/spacetx/starfish). Here, the images were normalized across channels and imaging cycles, a spot detection step was performed and, for each detected spot the intensity across all channels and cycles was extracted. The color sequence of each spot across cycles is matched to a decoding table that associates a color sequence with a specific gene. The output of this decoding is a csv table, in which each row represents a detection spot with different properties (x and y position, gene identity, quality metrics). The quality metrics for each spot is computed as follows: for each spot, the normalized fluorescence intensities across all channels are extracted. The prominent channel is considered the “true signal”, while all the others are considered “background”. The score is described by the formula “true signal” / (”true signal” + “background”), and it has a theoretical maximum of 1 (perfect decoding) and a theoretical minimum value of 0.25 when decoding in 4 colors, which corresponds to a random assignment (i.e. all the channels have the same fluorescence intensity for that spot). The quality score for each spot is computed per cycle, allowing to calculate 2 parameters: 1) the average quality across all cycles and, 2) the minimum quality across cycles. We found that filtering according to a minimum quality produces more reliable data, and we normally use a minimum quality value of 0.5.

#### Cell segmentation, clustering and EdU analysis

To identify cells, we used the DAPI staining of the nuclei. We applied the ’2D_versatile_fluo’ model from stardist (Schmidt et al., 2018) to the DAPI images, so to segment all the nuclei into ROIs. We manually checked the segmentation mask, to exclude the possibility of frequent segmentation artifacts. We then expanded each ROI of the mask, allowing each cell to “grow” isometrically for 20 pixels in each xy direction, unless the expansion clashed with a nearby expanding cell. We then assigned all the reads contained within the expanded ROIs to each cell, and save the resulting matrix into an AnnData object for downstream analysis. We then clustered the segmented cells using scanpy (Wolf et al., 2018), performing dimensionality reduction followed by Leiden clustering at different resolutions. We then critically assessed the spatial distribution of the inferred cell types under different clustering resolution, and when satisfied we extracted the most informative markers for each cluster, to assign it an identity based on available knowledge. We finally went back to the non-expanded DAPI mask and applied it onto the EdU image, extracted the intensity values for each pixel in every ROI, and computed the median EdU intensity for each ROI. We then filtered all the ROIs whose median was above an arbitrary threshold that labeled only the brightest EdU cells across the tissue, and labeled them in the AnnData object as EdU+ cells for downstream analysis.

## AUTHOR CONTRIBUTIONS

ERA, MG and FGM conceived the project, interpreted the data, designed and performed experiments and data analysis, and wrote the manuscript with input from all authors. ERA, AQ, AB and AMA performed single cell RNA sequencing. EV and RSG performed scRNAseq analysis. LE and ERA performed FACS sorting analysis. MG, SMS and ERA performed and analyzed the *in-situ sequencing* experiments. MN, AD, JME and FGM provided reagents. FGM led and coordinated the team.

## ACKOWLEDGEMENTS

The ISS unit at SciLifeLab provided some ISS imaging data.

## DATA AVAILABILITY

*Data will be made publicly available upon acceptance for publication*.

## SUPPLEMENTARY FIGURES AND FIGURE LEGENDS

**Figure S1.**
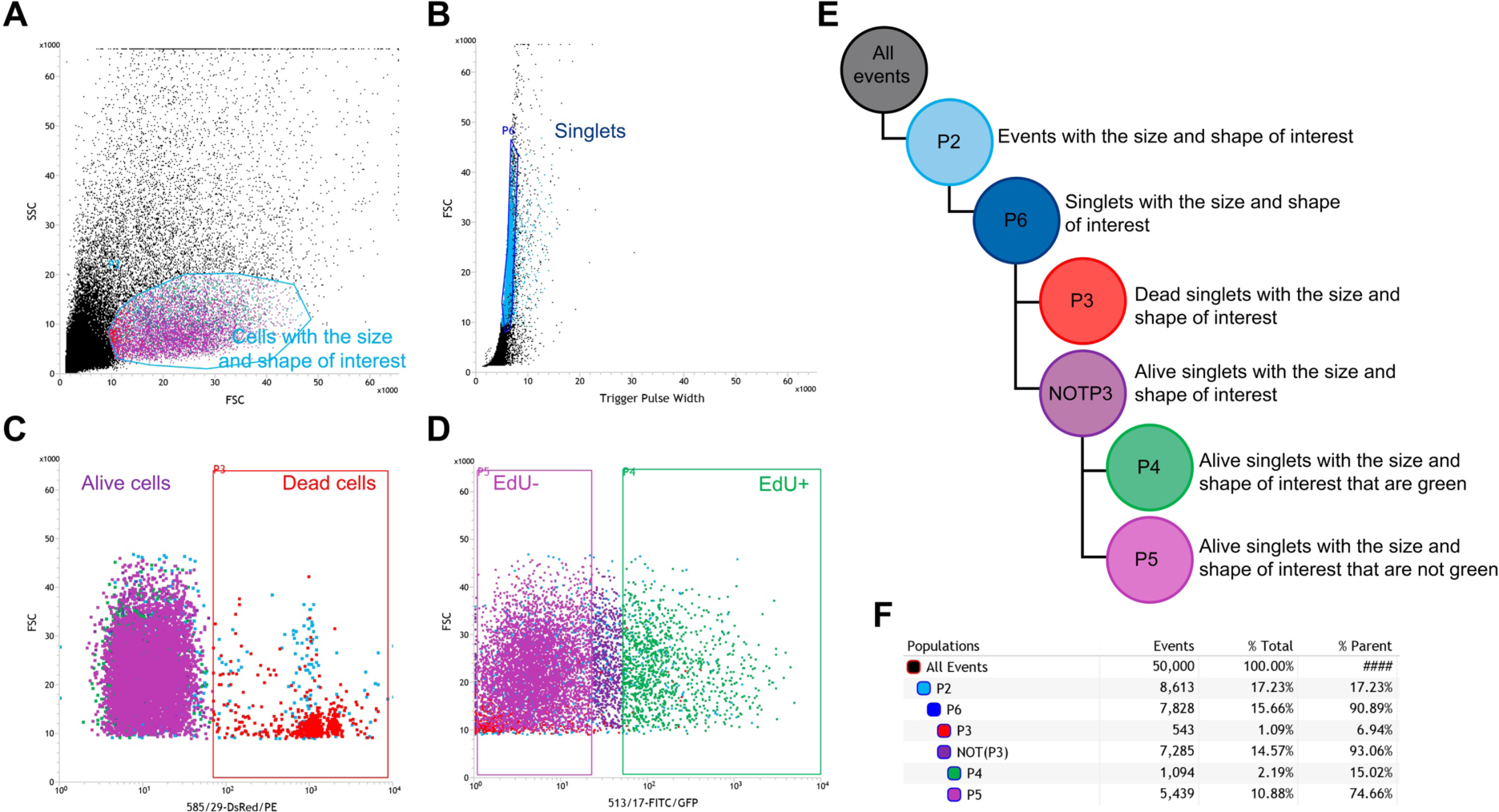
Graphic representation of the cell hierarchy defined for BirthSeq experiments. [A-D] Example of FC profiles obtained after the analysis of chick E14 neurons birthdated with EdU at E6. Each dotplot shows a total cell population (directly gated and identified in the dotplot). First, cell debris and nuclei are excluded and cells (P2, blue) with adequate size (high FCS) and shape (low SSC) are identified in SSC *versus* FSC dotplot [A]. Second, singlets (P6, dark blue) are gated in FCS *versus* trigger pulse width [B]. Third, dead cells (P3, red) are defined as PI^+^ in FSC *versus* dsRed dotplot [C]. Finally, EdU+ (P4, green) and EdU-(P5, purple) cells are identified as GFP^+^ and GFP^-^ in FSC *versus* GFP dotplot [D]. [E] Schematic representation of the defined cell hierarchy and parental populations. [F] Example of the statistical analysis obtained after a FC analysis. In the first column, cell hierarchy and all the gated populations are shown. In the events column, the number of events of each population is shown. Next, their relative proportions as total (third column) and parental (forth column) populations are shown.

**Figure S2.**
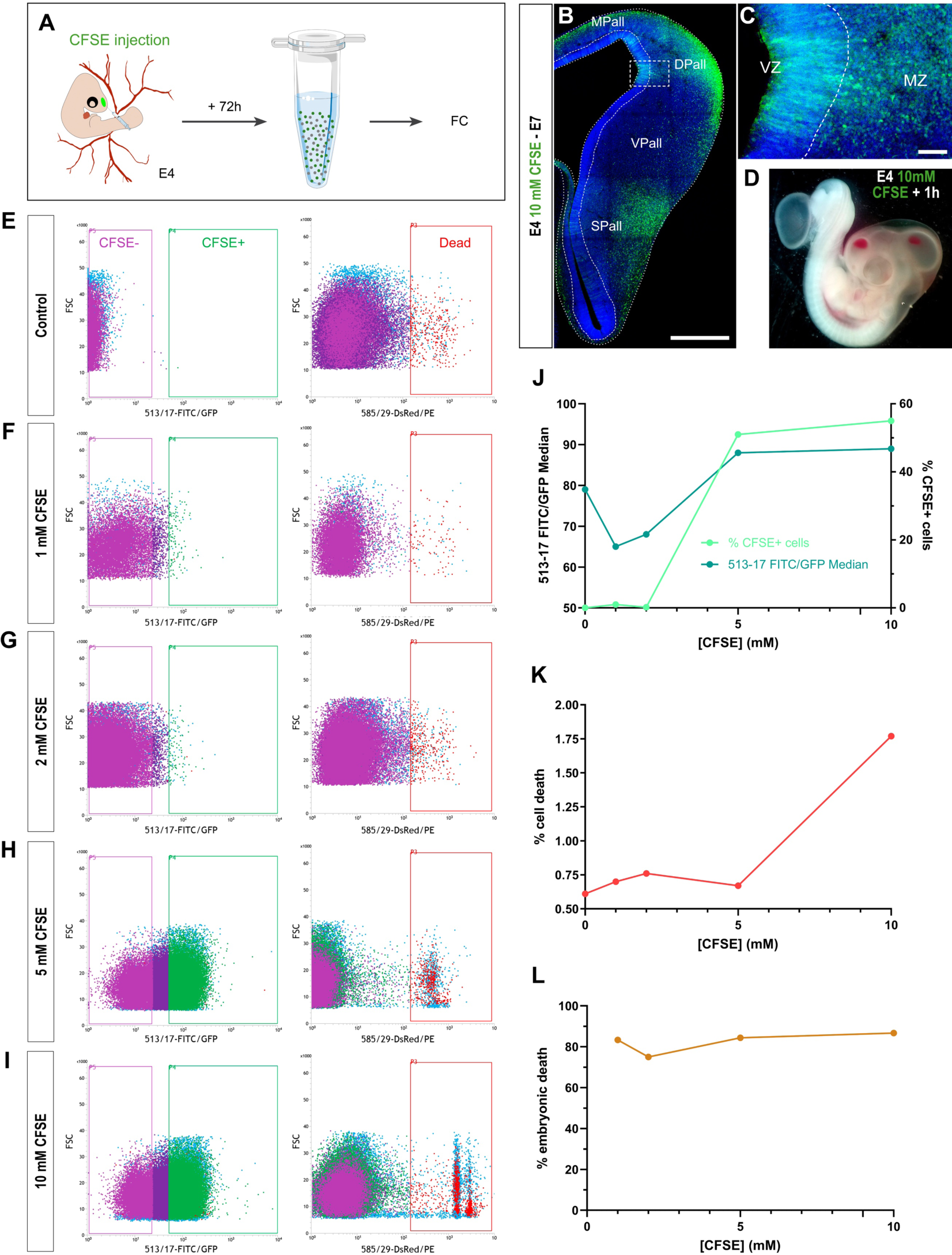
FlashTag is an ineffective birthdating system in chick embryos. [A] Experimental design of the FC analysis of chick embryonic neurons birthdated with CFSE. [B-C] Coronal section [B] and high-magnification view [C] of E7 chick telencephalon 3 days after 10 mM CFSE injection. It is possible to see CFSE+ cells both in the ventricular and in the mantle zone. [D] Representative picture of an E4 chick embryo with brain hemorrhages 1 hour after 10 mM CFSE injection. [E-I] FC profiles of chick E7 brain dissociated cells after 0 mM [E], 1 mM [F], 2 mM [G], 5 mM [H] and 10 mM [I] CFSE injections. [J-K] Graphic representation of FC analysis. Relative proportion of CFSE+ [J] and dead [K] cells were analyzed. CFSE+ cell brightness was assessed by 513-17 FITC/Median [J]. FC analysis revealed that high CFSE concentration (10 mM) is needed to be able to detect and isolate birthdated chick neurons, although is really damaging for the cells. [L] Analysis of chick embryonic death after 1 mM, 2 mM, 5 mM and 10 mM CFSE injections suggests that CFSE is toxic for chick embryos. *For nomenclature, refer to the list of abbreviations. DAPI counterstain in blue. Dashed white lines demarcate anatomical boundaries. Scale bars, 500 µm [B], and 50 µm [C]. CFSE+ (P4, green) and CFSE-(P5, purple) cells were identified as GFP^+^ and GFP^-^ in FSC versus GFP dotplot (left panel), whereas dead cells (P3, red) were defined as PI^+^ in FSC versus dsRed dotplot (right panel). Data represented in [J] and [K] graphs was obtained from a single cytometry experiment, and analyzed samples contained 3 pooled brains*.

**Figure S3.**
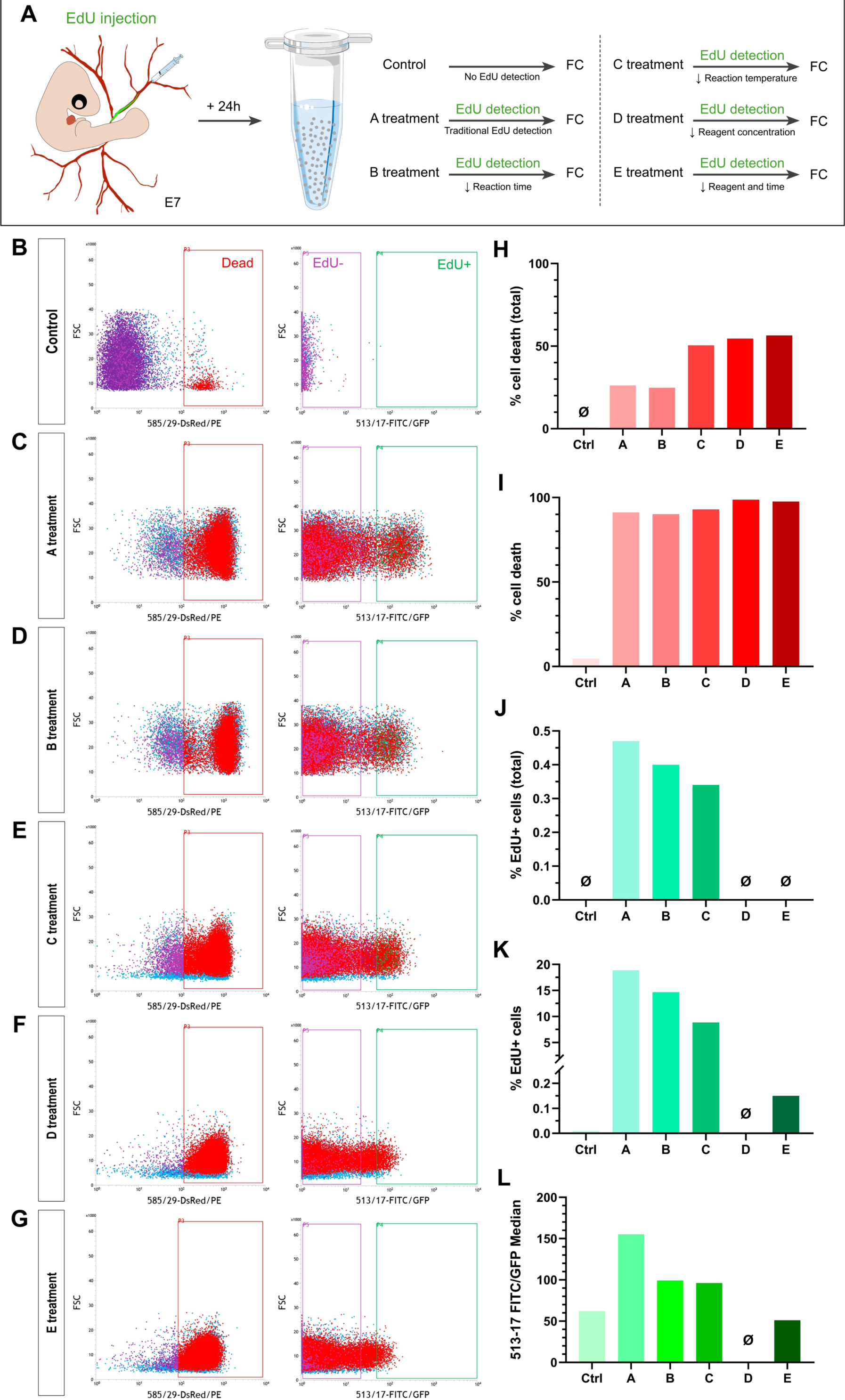
Modification of time, temperature and reagent concentration in the EdU detection process do not increase cell viability. [A] Experimental design of the FC analysis of dated chick embryonic neurons after time, temperature and reagent concentration variations of the EdU detection process. [B-G] FC profiles of dissociated chick embryonic neural cells 1 day after EdU injection and different EdU detection treatments: no EdU detection [B]; standard EdU detection protocol [C]; time [D], temperature [E] and reagent [F] modifications; both temperature and reagent [G] modifications. [H-L] Graphic representation of FC analysis. Relative proportion of dead [H-I] and EdU+ [J-K] cells of total and parental populations were analyzed. EdU+ cell brightness was assessed by 513-17 FITC/Median [L]. FC analysis showed that modifications of the EdU detection conditions do not only reduce EdU+ cell detection efficiency but also increase cell toxicity. *All the EdU detection protocol modifications are detailed in Methods. Dead cells (P3, red) were defined as PI^+^ in FSC versus dsRed dotplot (left panel) while EdU+ (P4, green) and EdU-(P5, purple) cells were identified as GFP^+^ and GFP^-^ in FSC versus GFP dotplot (right panel). Data represented in the graphs was obtained from a single cytometry experiment, and analyzed samples contained 3 pooled brains*.

**Figure S4.**
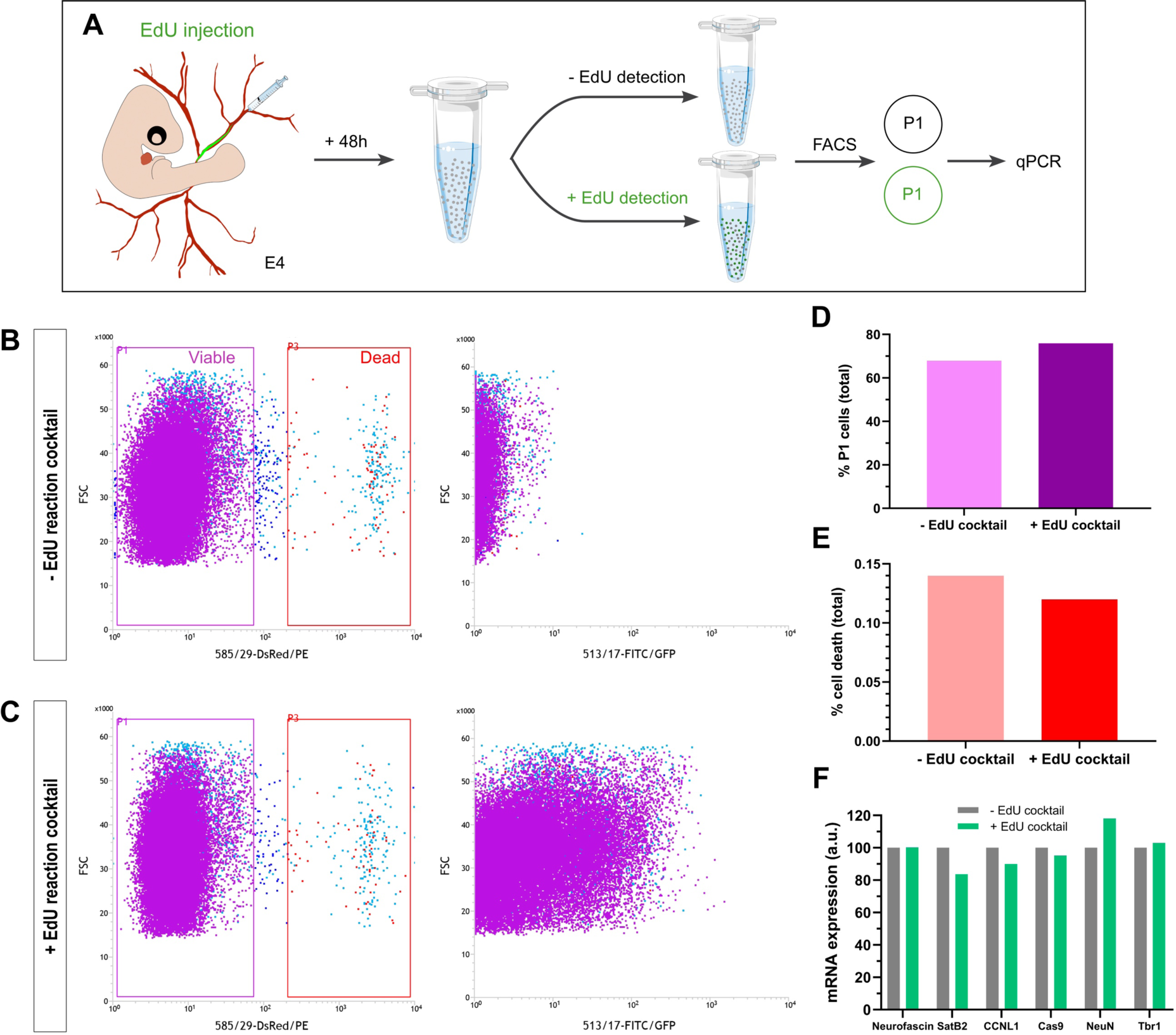
EdU detection cocktail does not affect gene expression. [A] Experimental design used to analyze gene expression after in revealed and non-revealed cells. [B-C] FC profiles of non-revealed [B] and revealed [C] chick embryonic neural cells 2 days after EdU injection. [D-E] Graphic representation of FC analysis. Relative proportion of viable [D] and dead [E] cells of total populations were analyzed. FC analysis showed that the treated sample contained GFP+ EdU+ cells, which were not detectable in the control sample. In addition, in both samples death population percentage was very low, ratifying the use of low Cu (II) containing detection cocktail in order to detect EdU+ cells without killing them. [F] Expression of neuronal (NEUROFASCIN, SATB2, NEUN and TBR1), cell division (CCNL1) and cell death (CAS9) genes in treated (+EdU cocktail) vs non-treated (-EdU cocktail) cells by RT-qPCR suggested that the detection cocktail was not altering gene expression. GAPDH was selected as a reference gene. *Viable (P1, purple) and dead (P3, red) cells were identified as PI^-^ and PI^+^ in FSC versus dsRed dotplot (left panel); whereas EdU- and EdU+ cells are shown in FCS vs GFP (right panel). Data represented in [D-F] was obtained from a single experiment. Each analyzed and sorted sample contained 3 pooled brains*

**Figure S5.**
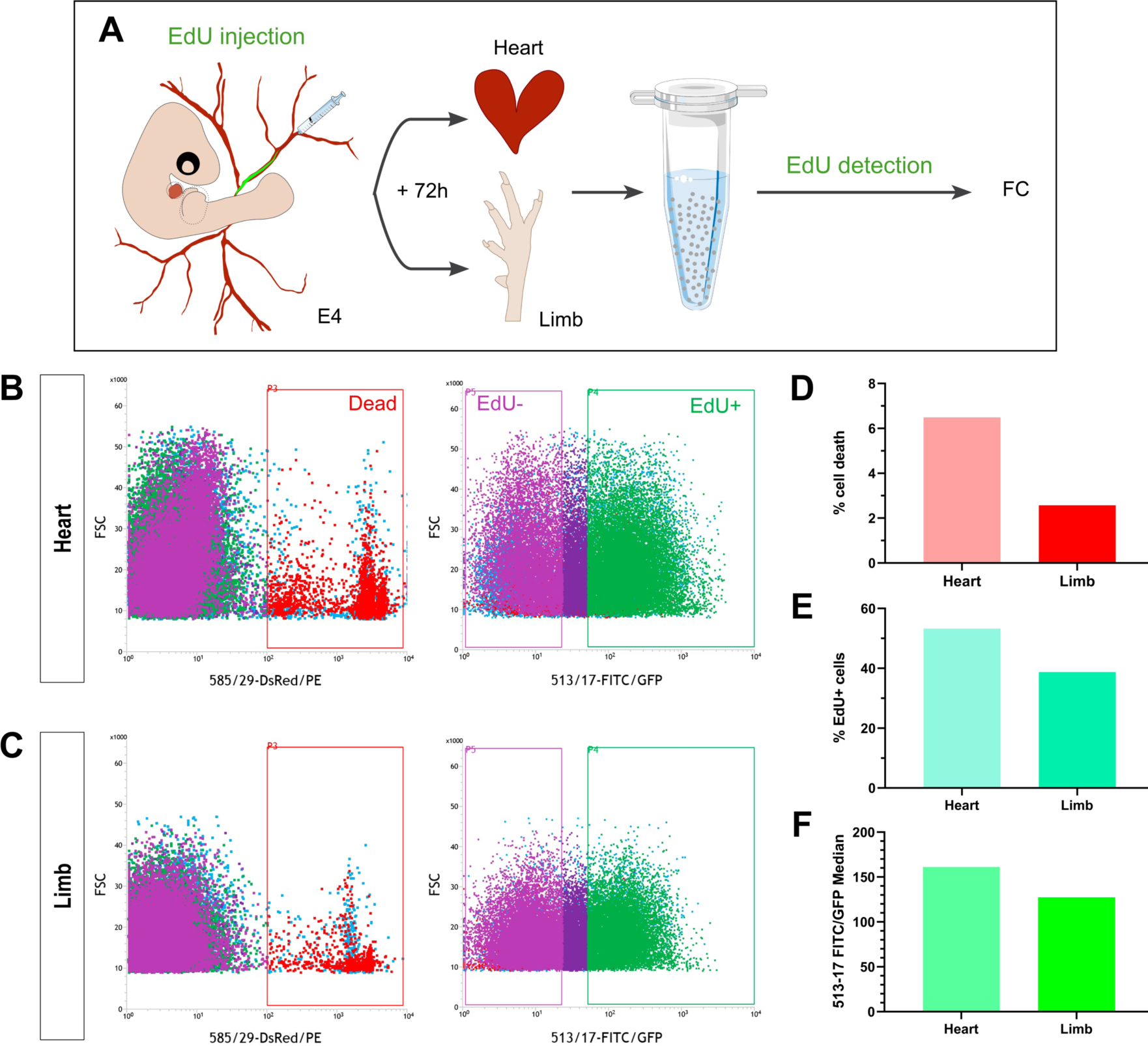
BirthSeq can be used to isolate birthdated cells from different organs. [A] Experimental design of the FC analysis of birthdated cardiac and limb chick embryonic cells. [B-C] FC profiles of dissociated cardiac [B] and limb [C] cells 3 days after EdU injection. [D-F] Graphic representation of FC analysis. Relative proportion of dead [D] and EdU+ [E] cells of parental populations were analyzed. EdU+ cell brightness was assessed by 513-17 FITC/Median [F]. FC analysis showed bright and live EdU+ populations in both limb and cardiac cell samples, demonstrating that this method works in different tissues. *Dead cells (P3, red) were defined as PI^+^ in FSC versus dsRed dotplot (left panel) whereas EdU+ (P4, green) and EdU-(P5, purple) cells were identified as GFP^+^ and GFP^-^ in FSC versus GFP dotplot (right panel). Data represented in the graphs was obtained from a single cytometry experiment. Heart sample contained 2 pooled E7 hearts, and limb sample contained 4 pooled limbs*.

**Figure S6.**
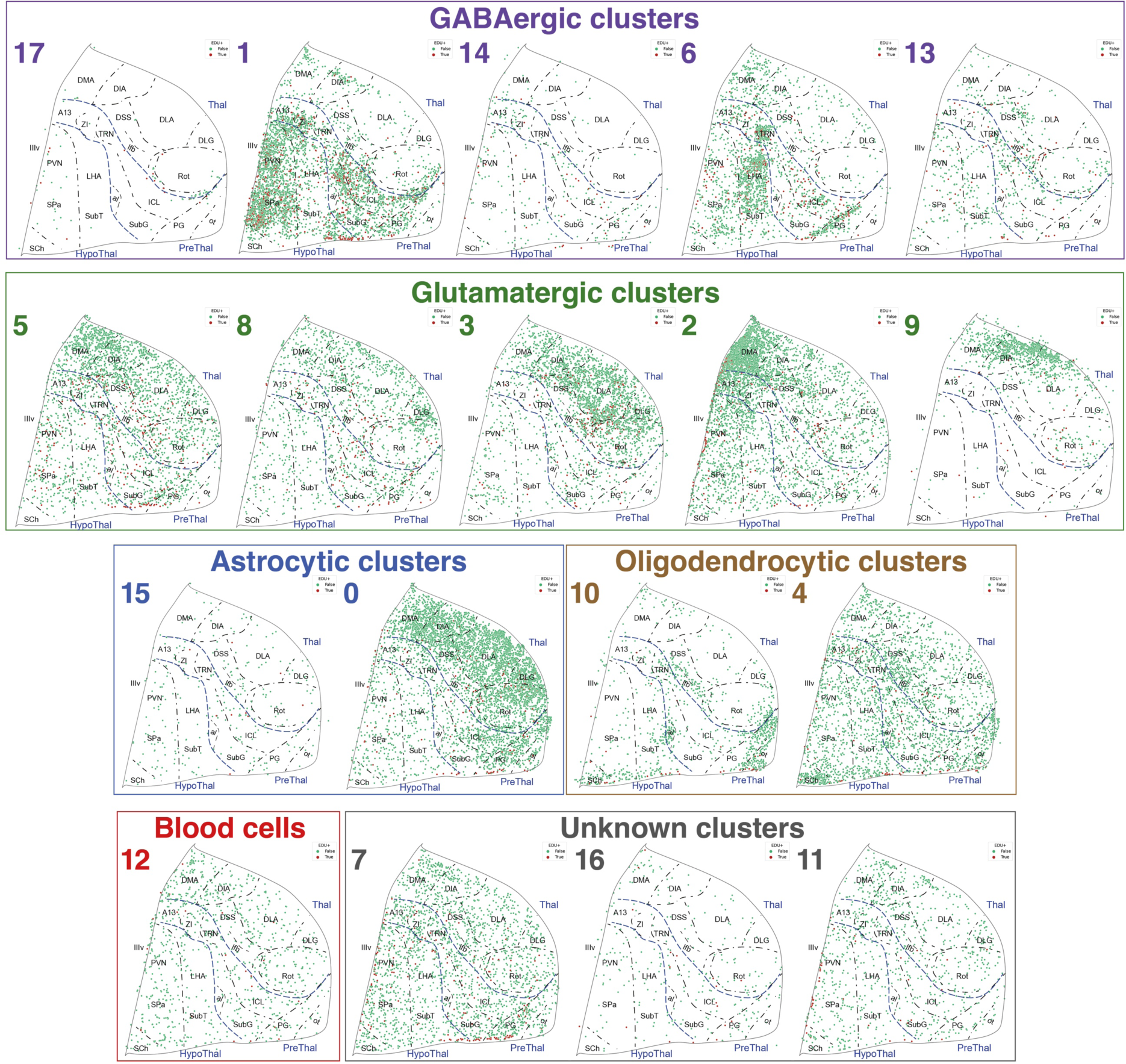
Diencephalic distribution of each cell cluster assigned by ISS and the location of E4-generated EdU+ neurons after NeurogenesISS. On the neuronal lineage, 5 different classes of GABAergic neurons and another 5 classes of glutamatergic neurons were identified. In the glial lineage, 2 types of astrocytes along with another two types of oligodendrocytes were differentially distributed within the diencephalon. In addition, we found one type of blood cells, and could not easily infer the cellular identity of three cell clusters.

